# Structural Adaptability in HIV-2 Vif Drives Broad Antagonism of APOBEC3G

**DOI:** 10.64898/2026.09.21.753272

**Authors:** Estelle K. Ronayne, Michelle Lilly, Xiaoxuan Lin, Yange Niu, Nicholas M. Chesarino, Michael Emerman, Yifan Cheng, John D. Gross

## Abstract

APOBEC3G (A3G), a restriction factor, is a barrier for zoonosis and targeted for degradation by the lentiviral protein Vif. Human Immunodeficiency Virus Type 1 (HIV-1) and HIV-2 arose from transmissions of lentiviruses from Old World monkeys to hominid primates. While HIV-1 Vif needed to adapt to specific residues in the arms race interface with A3G, HIV-2 Vif has greater tolerance for diversity there. Here, our cryo-EM structure of HIV-2 Vif and human A3G allows us to explore the mechanism underlying HIV-2 Vif’s broad specificity. While HIV-2 Vif does bind the arms race interface on A3G, HIV-2 Vif encodes tryptophans with low backbone stability to engage another interface on A3G. We propose a model where flexible loops in HIV-2 Vif undergo conformational selection to recognize diverse A3Gs, whereas the equivalent loops in HIV-1 Vif are preorganized to bind hA3G.

## Introduction

Human Immunodeficiency Virus Type 1 (HIV-1) and HIV-2 arose from zoonotic transmissions of Simian Immunodeficiency Viruses (SIVs) from natural reservoirs in Old World monkeys to hominid primates ^1^. APOBEC3G (A3G), a powerful restriction factor, acts as a barrier to zoonosis by blocking retroviral replication via catastrophic mutation to the viral genome ^2^. To prevent viral restriction, the lentiviral protein Vif binds and targets A3G for degradation by hijacking the host’s ubiquitin proteasome system. To do this, Vif assembles with host cofactors, CBFβ, EloB, and EloC, to form a substrate receptor (VCBC) for an E3 ligase which then ubiquitinates A3G ^3–5^.

Recent structural studies revealed that HIV-1 Vif binds human A3G (hA3G) via an RNA-mediated interface and a direct protein-protein interface ^6–8^. The RNA-mediated interface is evolutionarily constrained as A3G cytidine deaminase domain 1 (CDA1) must bind to viral RNA before A3G CDA2 can hyper-edit viral cDNA after reverse transcription ^2,6,9,10^. This places strong evolutionary pressure on the direct protein-protein interface, also known as the arms race interface, which includes Vif loop 5 and position 128 of A3G ^6–8^.

A3G and Vif are involved in an ancient host-virus arms race, where A3G evolves to escape Vif binding and Vif adapts to re-establish complex formation ^11–13^. Across primate species, there is sequence variation in A3G at position 128 that creates a barrier for zoonosis of primate lentiviruses ^11,13^. The transmission of SIV from red-capped mangabey monkeys (SIVrcm), a precursor to HIV-1, into hominids required adaptation in Vif loop 5 (Y86 to H/Q, equivalent to HIV-1 Vif Q83) for robust neutralization of hominid A3G, which encodes D128 ^12,14^. Similarly, HIV-1 Vif can successfully degrade hA3G but is unable to neutralize A3Gs that encode basic residues at position 128, such as rcmA3G which encodes K128 ^15,16^.

In contrast, HIV-2 Vif can broadly neutralize A3G with diverse identities at position 128 ^12,17^. Additionally, both HIV-2 and its precursor, SIV from sooty mangabey monkeys (SIVsmm), encode a Vif protein with a tyrosine in loop 5 (HIV-2 Vif Y85, equivalent to SIVrcm Vif Y86) and can successfully neutralize hA3G, indicating a Y to H/Q adaptation in Vif loop 5 was not required for the HIV-2 lineage ^1^. However, the reasons why HIV-2 Vif can antagonize A3Gs with diverse sequences at the arms race interface, or why adaptation in HIV-2 Vif loop 5 was not necessary, remains a puzzle.

Here, by determining a cryo-EM structure of hA3G bound to HIV-2 Vif and its host cofactors, CBFβ, EloB, and EloC (HIV-2 VCBC), we provide further insights into how HIV-2 Vif can accommodate diverse sequences at the arms race interface. While the overall structural architectures of HIV-1 and HIV-2 Vif in complex with hA3G are very similar, sequence variation in Vif loop 3 (Trp-Rich loop in HIV-2) and loop 5 (the molecular arms race loop) results in conformational and energetic differences. At both interfaces, HIV-2 Vif encodes tryptophan residues with low backbone stability which directly engage hA3G through side chains interactions. As a result, HIV-2 Vif engages more total buried solvent accessible surface area (BSASA) at the protein-protein interface with hA3G than HIV-1 Vif. We propose these loops in HIV-2 Vif undergo conformational selection to allow recognition of diverse A3Gs, whereas the equivalent loops in HIV-1 Vif are preorganized to bind hA3G resulting in more narrow specificity.

## Results

### Cryo-EM Structure of Human A3G, HIV-2 Vif, and Host Co-factors

While previous HIV-2 Vif studies have focused on Vif from the laboratory adapted strain HIV-2_ROD_, we screened several full-length primary isolates of HIV-2 Vif sequences for our study ^17–19^. After testing several candidates for co-expression with hA3G, CBFβ, EloB, EloC, and the N-terminus of Cullin 5 (residues 1-386, Cul5N) in insect cells, we selected Vif from the HIV-2_ALI_ strain for further study because it expressed well in insect cells and exhibits strong neutralization of hA3G that is comparable to HIV-2_ROD_ Vif (**Figure** S1A-B). We determined a 3.6Å cryo-EM structure of full length, wild type (WT) hA3G and HIV-2 VCBC (**Figure** 1A-B, S1-3, Table 1). While Cul5N is co-purified with the complex, there is no cryo-EM density for the Cul5N component due to either intrinsic flexibility or dissociation during freezing (**Figure** S1C-D). HIV-2 Vif and hA3G interact via RNA-mediated, arms race, and Trp-Rich interfaces (**Figure** 1C-E).

**Figure 1.**
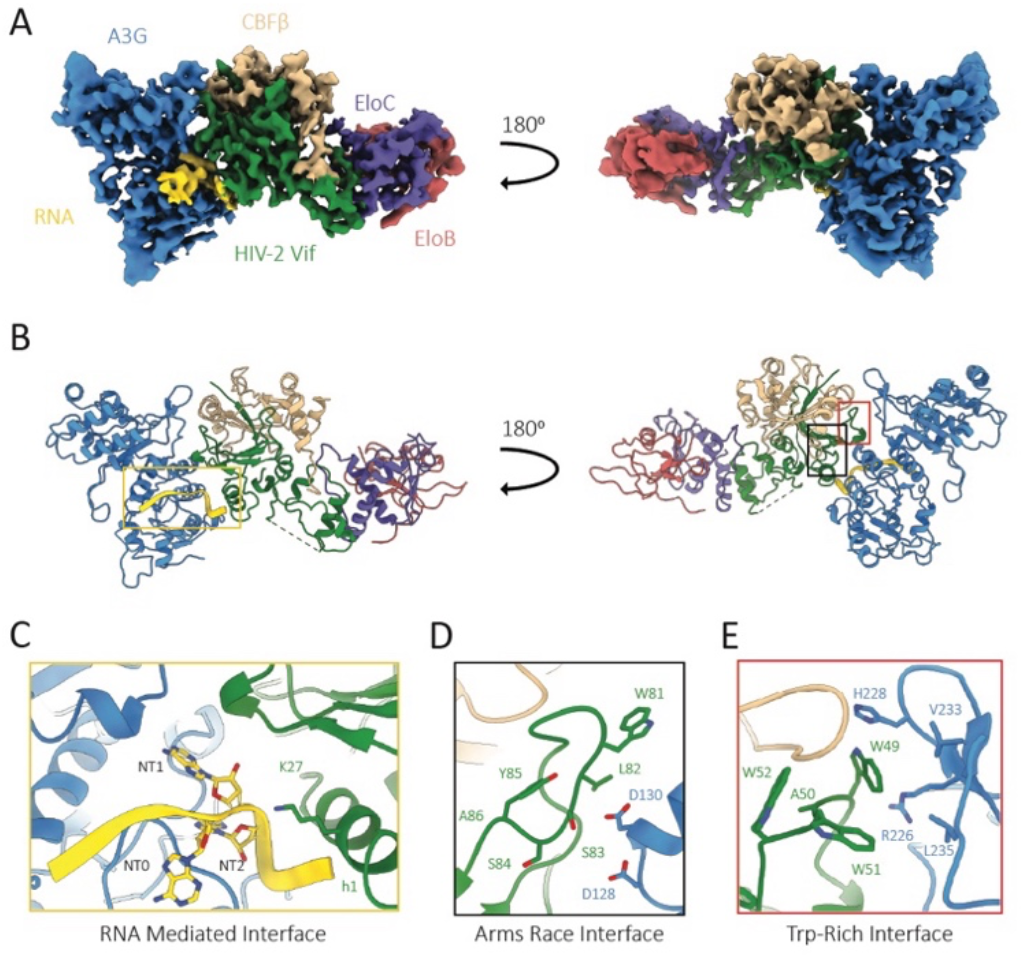
Cryo-EM structure of human A3G, HIV-2 Vif, and host co-factors. **A)** Sharpened cryo-EM map for the hA3G and HIV-2 VCBC complex. **B)** Atomic model of the hA3G and HIV-2 VCBC complex with boxes indicating the three interfaces between hA3G and HIV-2 Vif. Here and throughout, HIV-2 Vif is shown in green, hA3G in blue, CBFβ in beige, EloB in pink, EloC in purple, and RNA in yellow. **C)** Single-stranded RNA mediated interface between hA3G CDA1 and HIV-2 Vif h1. **D)** Arms race interface between HIV-2 Vif loop 5 and positions 128 and 130 of hA3G. **E)** Trp-Rich interface between HIV-2 Vif loop 3 and hA3G CDA2.

Our HIV-2 VCBC and hA3G structure shares a similar quaternary structure to previous HIV-1 structures ^6–8^. As in the HIV-1 Vif and hA3G structures, a single-stranded RNA-mediated interface is present with a four nucleotide (NT1-4) core acting as a molecular glue between A3G CDA1 and HIV-2 Vif helices 1 and 3_10_ (**Figure** 1C). The overall folds of HIV-1 and HIV-2 Vif are similar with a Root Mean Square Deviation (RMSD) of 2.9Å across all atoms and 1.1Å for the structurally conserved core, with notable differences in loop 5 conformation and a shorter helix 1 in HIV-2 Vif (**Figure** S4A-B) ^6^. Additionally, there is a longer loop connecting helices 4 and 5 in HIV-2 Vif, most of which remains unresolved (residues 161-200). The hA3G structures in the HIV-1 and HIV-2 complexes are overall in high agreement with a RMSD of 0.7Å (**Figure** S4C-D). At the Trp-Rich interface, HIV-2 Vif loop 3 (Trp-Rich loop) encodes 48-GWAWW-52 and packs against a beta hairpin of hA3G, with A3G V233 and L235 (**Figure** 1E). At this interface, HIV-2 Vif W49 and W51 are buried at the HIV-2 Vif and hA3G interface, while W52 is nestled against CBFβ. Additionally, hA3G R226 is positioned to form a cation-pi interaction with HIV-2 Vif W49. Notably, at the Trp-Rich interface of the HIV-2 complex, the beta hairpin encoding hA3G H228 is rearranged and results in a closer contact to HIV-2 than HIV-1 Vif (**Figure** S4C) ^6^. Near the Trp-Rich interface, CBFβ loop 6 co-folds differently with HIV-2 than HIV-1 Vif (**Figure** S5A-B). In our structure, CBFβ loop 6 lays against HIV-2 Vif loop 3 and the beta sheet core, as in the SIVrcm VCBC complex, whereas CBFβ contacts HIV-1 Vif loop 3 but does not engage the beta sheet core as extensively (**Figure** S5) ^6,14^. Despite the overall similarity, there is about 80Å^2^ more total BSASA at the protein-protein interface between hA3G and HIV-2 Vif (641Å^2^) than with HIV-1 Vif (560Å^2^) due to increased interactions at the arms race and Trp-Rich interfaces. As HIV-2 Vif loop 5 engages more surface area on hA3G, we speculated this larger interface at the canonical arms race may underlie HIV-2 Vif’s broad specificity.

### HIV-2 Vif Loop 5 Does Not Confer HIV-1 Vif with Broad Antagonism of A3G

While loop 5 of both HIV-1 and HIV-2 Vif bind the arms race interface on hA3G, only the HIV-1 lineage required adaptation in Vif loop 5 (SIVrcm Vif Y86 to H/Q) for cross-species transmission into hominids. Vifs in the HIV-2 lineage can broadly neutralize A3G without an equivalent adaptation. Notably, loop 5 of HIV-1 and HIV-2 Vif adopt different conformational states to recognize hA3G (**Figure** 2A-B). The adaptive residue in HIV-1 Vif, Q83 (equivalent to SIVrcm Vif Y86 and HIV-2 Y85), directly binds the arms race interface of hA3G and is well resolved, but most of HIV-1 Vif loop 5 does not contact hA3G and remains poorly resolved (**Figure** 2B) ^6^. In contrast, HIV-2 Vif loop 5 makes more extensive contacts with hA3G but HIV-2 Vif Y85 does not appear to directly bind the arms race interface of hA3G. As the local resolution of HIV-2 Vif loop 5 is relatively low, it is difficult to determine many sidechain rotamers at the arms race interface (**Figure** S3D). However, our cryo-EM map enables confident modeling for most of the HIV-2 Vif loop 5 backbone. Together, these results demonstrate HIV-1 and HIV-2 Vif loop 5 engage the arms race interface on hA3G differently.

To assess whether the additional contacts between HIV-2 Vif loop 5 and hA3G help impart tolerance for variety at the arms race interface, we performed partial and full loop 5 swaps between HIV-1 and HIV-2 Vif, with the well conserved tryptophan in Vif loop 5 as a boundary for our partial swaps. Then we tested these Vif mutants against hA3G with both the wild-type sequence at position 128 (D128) and the amino acid at position 128 most frequently found in Old World monkey A3G (K128) (**Figure** 2C-D, S6A-B). We observed an overall reduction in infectivity for these mutants, particularly for the full loop 5 swaps. To our surprise, we found that HIV-1 Vif with all or part of HIV-2 Vif loop 5 maintained its sensitivity to position 128 of hA3G (**Figure** 2D). Similarly, HIV-2 Vif with all or part of HIV-1 Vif loop 5 maintained relative insensitivity to substitution at position 128 of hA3G, with HIV-2 Vif Full and Partial 1 swaps displaying slightly increased infectivity against D128K hA3G compared to WT hA3G. Therefore, we conclude that determinants outside of HIV-2 Vif loop 5 contribute to HIV-2 Vif’s broad neutralization of A3G.

**Figure 2.**
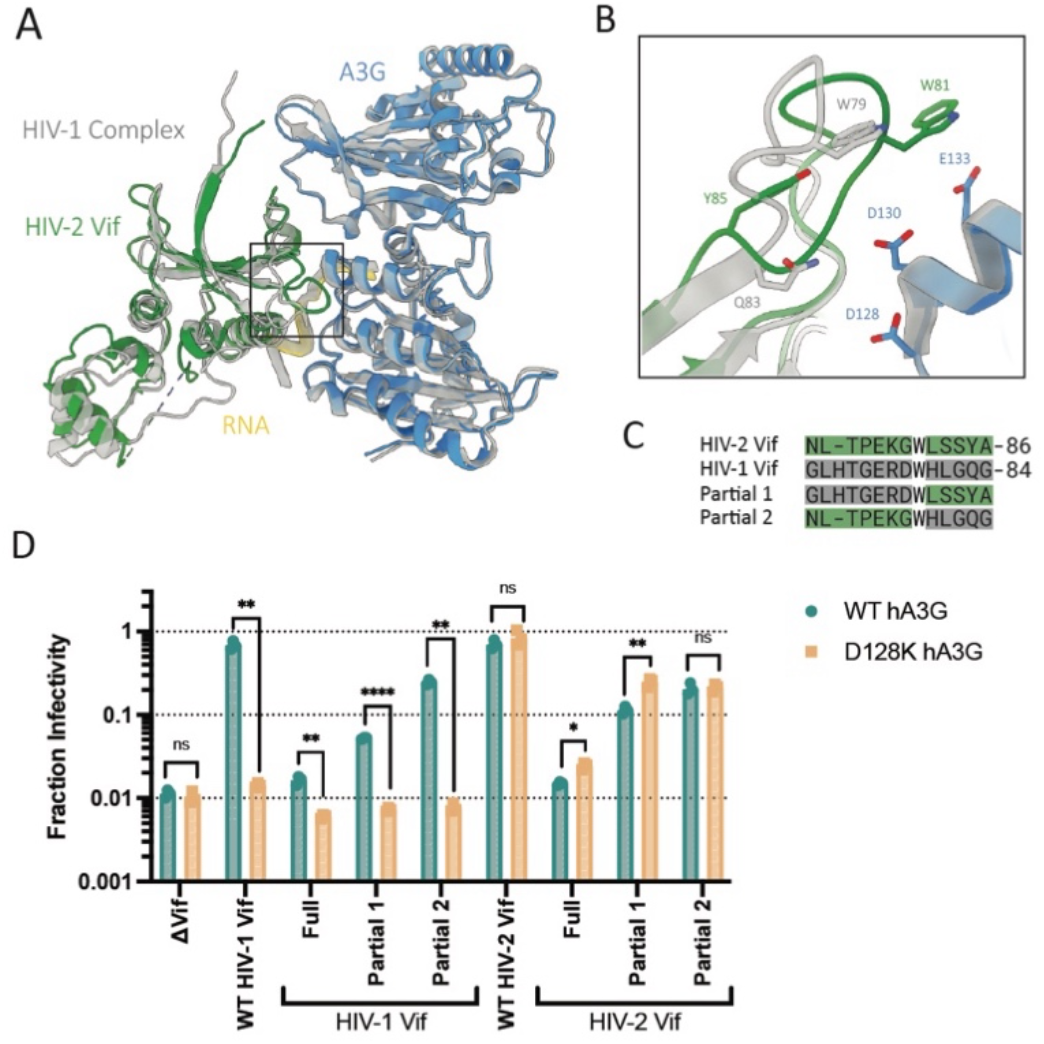
HIV-2 Vif loop 5 interacts differently with hA3G than that of HIV-1 Vif. **A)** HIV-2 Vif and hA3G complex (colored) overlayed with the HIV-1 Vif and hA3G complex (grey; PDB: 8CX0), aligned by hA3G. **B)** Close-up of the arms race interface with HIV-2 Vif Y85 and W81, equivalent residues in HIV-1 Vif, and positions 128 and 130 of hA3G shown as sticks. **C)** Loop 5 sequences of HIV-2 Vif, HIV-1 Vif, and the partial swaps. **D)** Single-cycle viral infectivity assay to assess the effect of the HIV-1 and HIV-2 Vif loop 5 swaps in the presence of WT hA3G (teal) or D128K hA3G (yellow). The Y-axis represents the fraction infectivity. Standard deviation from the mean of three biological replicates is represented by error bars. * P < 0.05, ** P < 0.01, *** P < 0.001, **** P < 0.0001 as calculated by the unpaired Welch’s t-test.

### HIV-2 Vif Trp-Rich Loop Forms an Additional Interface with hA3G

We hypothesized that HIV-2 Vif may accommodate for variation at the arms race of A3G due to additional protein-protein interactions elsewhere. Thus, we examined the Trp-Rich interface between HIV-2 Vif and hA3G (**Figure** 3A-B). The interface between HIV-2 Vif loop 3 and hA3G (199Å^2^) is over 150Å^2^ larger than the equivalent interface in the HIV-1 Vif and hA3G complex (46Å^2^). Given the structural significance of this unique interface, we reasoned that the HIV-2 Vif Trp-Rich loop could be responsible for HIV-2 Vif’s broad specificity of A3G. Among HIV-2 Groups A and B, the “GWAWW” sequence in Vif loop 3 is strongly conserved (**Figure** 3C). There is more variation in Vif loop 3 across HIV-1 Group M, but none contain the same sequence as HIV-2 in this region (**Figure** 3D). Swapping Vif loop 3 between HIV-1_LAI_ Vif (PHPR) and HIV-2 Vif (GWAWW) resulted in a strong loss of function for both mutants, with fractions of infectivity near the ΔVif controls (**Figure** S7). While these Vif loop 3 swaps suggest the loop is functionally critical, the infectivity levels of these mutants were not high enough to determine the differential infectivity against WT and D128K hA3G.

Therefore, to study the importance of specific residues comprising this Trp-rich loop in HIV-2 Vif, we turned to SIVrcm Vif which encodes a similar Trp-Rich motif but does not have broad tolerance for variation at the arms race of A3G (**Figure** 3C and E). When comparing Vif loop 3 sequences, we noted the residues preceding and spacing the Trp-Rich motif are different in HIV-2 Vif and SIVrcm Vif. Interestingly, the Trp-Rich motif is preceded by a glycine in HIV-2 Vif and a proline in SIVrcm Vif. Given glycine and proline can access the most and least dihedral angles respectively, it is likely that a glycine residue would allow more backbone flexibility in the Vif Trp-Rich motif than a proline. Thus, we predicted the backbone flexibility of Vif’s Trp-Rich motif would modulate antagonism of A3G.

### HIV-2 Vif Encodes Backbone Flexibility at the hA3G Binding Interface

To investigate how backbone flexibility of Vif loop 3 affects antagonism of A3G, we performed Hydrogen-Deuterium eXchange Mass Spectrometry (HDX-MS) experiments on HIV-1, HIV-2 and SIVrcm Vif bound to cognate A3G and host factors (**Figure** 4, S8-9, Table 2). HDX-MS measures the exchange rate of backbone amide hydrogens with the solvent deuterium. Exchange occurred via EX2 kinetics, in which the observed HDX rates reported on the equilibrium between the closed, H-bonded state of backbone amide and its transiently open, unfolded state that is exchange-competent. Hence, HDX-MS directly reports on local stability, providing a measurement of the folding free energy (ΔG) that can be traced to residues of interest, where ΔG values closer to zero indicate less stable, more dynamic regions whose backbone H-bonds undergo more frequent transient breakage.

**Figure 3.**
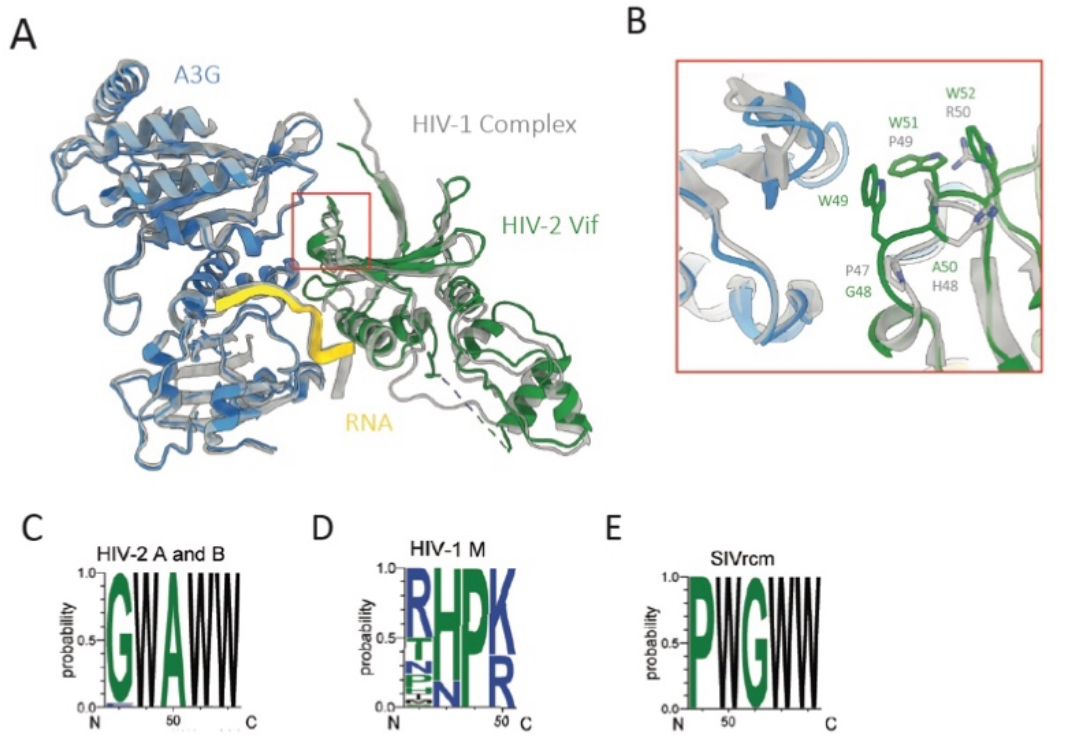
HIV-2 Vif Trp-Rich loop, not present in HIV-1 Vif, forms an additional interface with hA3G. **A)** HIV-2 Vif and hA3G complex (colored) overlayed with the HIV-1 Vif and hA3G complex (grey; PDB: 8CX0), aligned by hA3G. **B)** Close-up of Trp-Rich interface with HIV-2 and HIV-1 Vif loop 3 residues shown as sticks. **C-E)** Logo plot of sequence variation of Vif loop 3 for **C)** HIV-2 Groups A and B (104 sequences), **D)** HIV-1 Group M (42 sequences) and **E)** SIVrcm (4 sequences).

Overall, loop 3 in HIV-2 Vif appears to have the lowest backbone stability, measured as the fastest HDX rates, followed by SIVrcm and then HIV-1 Vif (**Figure** 4A-C, S8). HIV-2 Vif also has the widest range of stability across loop 3 residues when compared to the other Vif species. HIV-2 W49, which contacts hA3G, and HIV-2 A50 showed the fastest H-D exchange, while HIV-2 W52, which contacts CBFβ, has the slowest exchange rate (**Figure** 4A). HIV-2 Vif G48 likely allows greater range of conformational sampling for the backbone of HIV-2 Vif W49, due to the wide range of permitted Ramachandran angles of glycine, while SIVrcm Vif is restricted by a proline residue at the equivalent position (**Figure** 4A-B). The conformational restriction by this proline in SIVrcm Vif may be compensated by the glycine spacing the Trp-Rich motif which likely accounts for the low backbone stability of SIVrcm Vif 50-WGW-52 (**Figure** 4B). It is unsurprising HIV-1 Vif loop 3 undergoes the least exchange since it is proline rich (PHPR) and forms a tight hairpin for the beta sheet (**Figure** 4C). Interestingly, we found that the region of hA3G at this interface exhibited slower HDX rates when in complex with HIV-2 Vif than HIV-1 Vif, suggesting a tighter binding interface between hA3G and HIV-2 Vif involving a greater binding energy (**Figure** 4D-F). These energetic profiles demonstrate that the Trp-Rich loop in HIV-2 Vif forfeits backbone stability to permit sufficient conformational sampling which enables extensive interactions between the tryptophan sidechains and hA3G.

**Figure 4.**
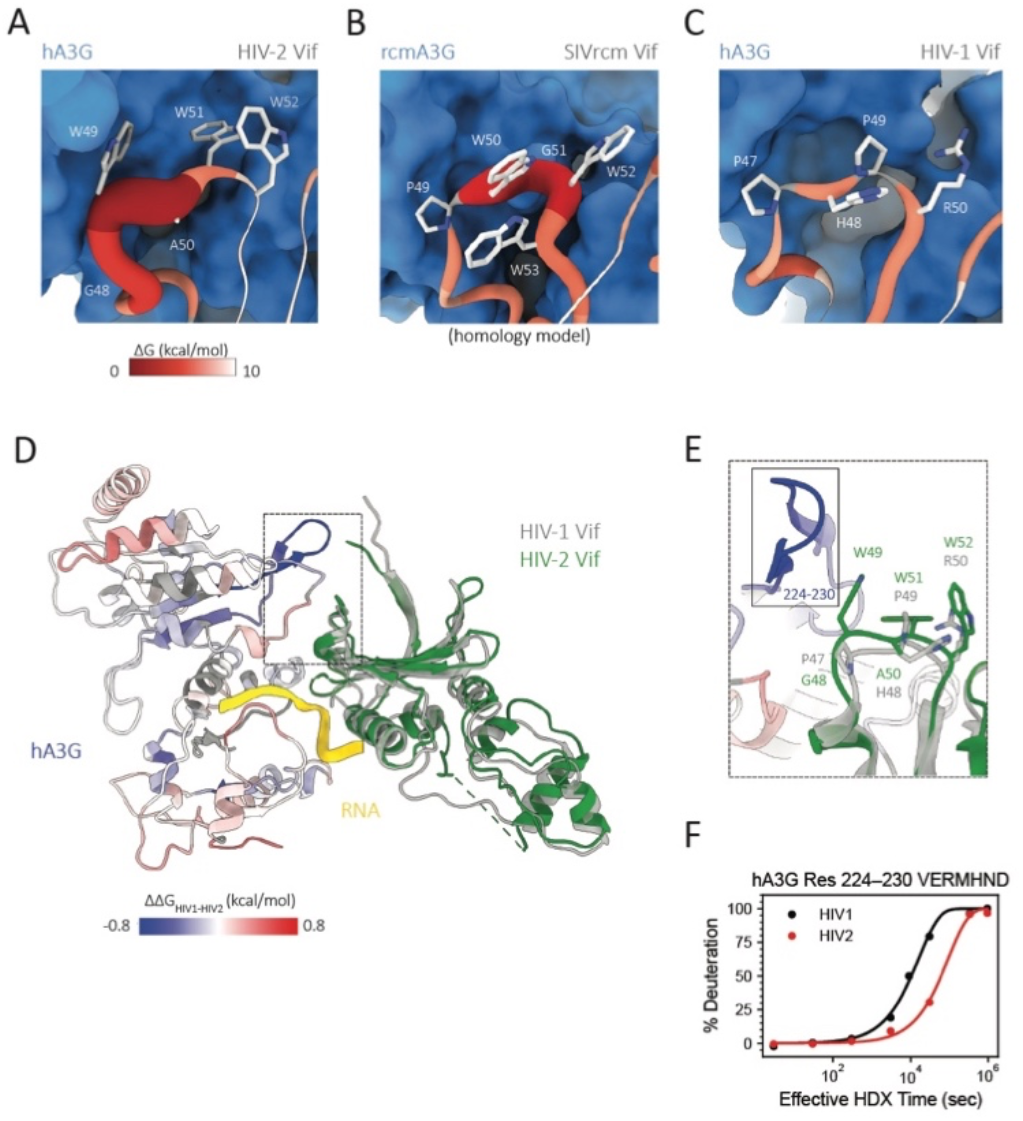
HIV-2 Vif encodes residues with low backbone stability at the Trp-Rich interface to engage hA3G. **A-C)** The Gibbs free energy of unfolding (ΔG) for **A)** HIV-2 Vif loop 3 while bound to hA3G, **B)** SIVrcm Vif loop 3 while bound to rcmA3G (visualized on a homology model of the rcmA3G and SIVrcm VCBC complex), **C)** HIV-1 Vif loop 3 while bound to hA3G (PDBID: 8CX0). A3G is shown as blue surface. Vif loop 3 is shown in grey sticks with backbone color and thickness reflecting ΔG values: red and thick for ΔG of 0 kcal/mol, grading to white and thin for ΔG of 10 kcal/mol. **D-E)** Difference in unfolding free energy of hA3G bound to HIV-2 Vif relative to HIV-1 Vif (ΔΔG_HIV1-HIV2_) in kcal/mol, with **E)** a closeup of the Trp-Rich interface. **F)** Kinetics of deuterium uptake for peptide 224-230 of hA3G at the binding interface.

At both the arms race and Trp-Rich interfaces, HIV-2 Vif encodes tryptophans with low backbone stability that directly engage hA3G (**Figure** 4A and S9A). Our energetic and structural analysis demonstrates that these residues in HIV-2 Vif adopt energetically unfavorable, low stability, backbone conformations in to form favorable sidechain interactions with hA3G. We propose a model where flexible loops in HIV-2 Vif undergo conformational selection to recognize diverse A3Gs, whereas the equivalent loops in SIVrcm and HIV-1 Vif occupy a more limited conformational space (**Figure** 4 and S9).

### Conformational Landscape of Vif Trp-Rich Loop Modulates Infectivity Against hA3G

Based on our HDX analysis, we predict Vif will form more extensive interactions with A3G, and therefore have greater tolerance for variation at the arms race interface, when a glycine precedes its Trp-Rich motif. Indeed, when replacing the proline in SIVrcm Vif with a glycine (SIVrcm P49G), we observe a modest increase in infectivity against both WT and D128K hA3G (fraction infectivity of 0.072 to 0.135 and 0.320 to 0.454, respectively), consistent with our energetic analysis (**Figure** 5A). Conversely, when the glycine spacing the Trp-Rich motif is replaced with an alanine (SIVrcm Vif G51A), which likely restricts the loop conformational landscape, infectivity against WT and D128K hA3G are reduced by about 14 and 6-fold, respectively (fraction infectivity of 0.072 to 0.005 and 0.320 to 0.057, respectively), suggesting weaker recognition of hA3G (**Figure** 5A). Strikingly, wildtype infectivity levels are rescued when the residues preceding and spacing the Trp-Rich motif in SIVrcm Vif are both replaced with their equivalents in HIV-2 Vif (SIVrcm P49G/G51A) (**Figure** 5A). Together, these results suggest SIVrcm Vif loop 3 plays a role in recognition of hA3G.

When the glycine preceding the Trp-Rich motif in HIV-2 Vif is replaced with a proline (HIV-2 Vif G48P), we observe a modest loss of function against WT hA3G and no effect on infectivity against D128K hA3G (**Figure** 5B). As with our energetic analysis and SIVrcm Vif infectivity assays, this data supports our model where increased conformational restraints in the Vif Trp-Rich loop reduce antagonism of hA3G and narrow Vif’s specificity for A3G. Curiously, HIV-2_ROD_ Vif G48A was previously reported to have a strong loss of function against hA3G, and this discrepancy may be due to the different conformational restraints imposed by glycine, proline, and alanine ^17^. There is no effect on infectivity against WT or D128K hA3G when the alanine spacing the Trp-Rich motif in HIV-2 Vif is replaced with glycine (HIV-2 Vif A50G) (**Figure** 5B). This suggests a glycine preceding HIV-2 Vif’s Trp-Rich motif allows sufficient sampling of backbone conformations for hA3G binding and any further increase in loop flexibility has no functional benefit. When the residues preceding and spacing the Trp-Rich motif in HIV-2 Vif are both replaced with their equivalents in SIVrcm Vif (HIV-2 Vif G48P/A50G), we observe a modest decrease in infectivity against WT hA3G comparable to the single glycine to proline substitution (**Figure** 5B). This demonstrates the conformational restraints imposed when a proline precedes the Trp-Rich motif in HIV-2 Vif are not alleviated by a flexible glycine spacing the motif. All the HIV-2 and SIVrcm Vif loop 3 mutants tested maintained full infectivity against rcmA3G, with HIV-2 Vif A50G having a modest increase in infectivity (**Figure** 5A-B). Thus, the reduced infectivity levels we observe for SIVrcm Vif G51A, HIV-2 Vif G48P and HIV-2 Vif G48P/A50G against hA3G reflect a fine-tuning of function rather than a general loss of A3G recognition. Together, these results demonstrate the subtle changes to the conformational landscape of the Vif Trp-Rich loop tune antagonism of hA3G.

**Figure 5.**
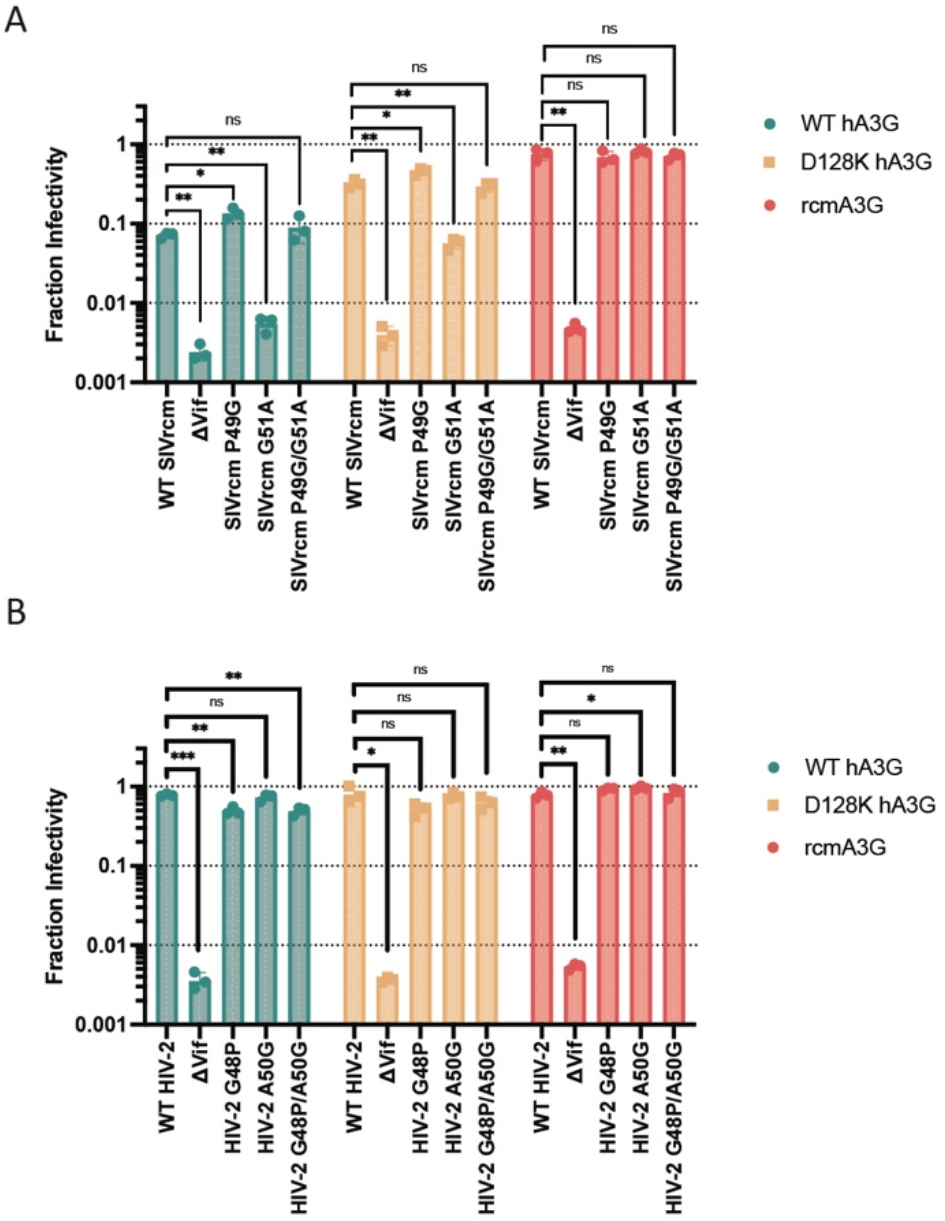
Preceding and spacing residues of Vif’s Trp-Rich loop modulate infectivity against hA3G. **A-B)** Single-cycle viral infectivity assay to assess the effect of mutation to non-tryptophan residues in Vif loop 3, with SIVrcm in **A** and HIV-2 in **B**, in the presence of WT hA3G (teal), D128K hA3G (yellow), or rcmA3G (red). The Y-axis represents the fraction infectivity. Standard deviation from the mean of three biological replicates is represented by error bars. * P < 0.05, ** P < 0.01, *** P < 0.001, **** P < 0.0001 as calculated by the unpaired Welch’s t-test.

## Discussion

While both HIV-1 and HIV-2 Vif loop 5 engage the arms race interface on hA3G directly, only HIV-2 Vif can broadly neutralize A3Gs with sequence variation at position 128 ^6–8,12,15–17^ (**Figure** 2A-B). Our study identifies an additional interface between a flexible Trp-Rich motif in HIV-2 Vif, which is not present in HIV-1 Vif, and another site on hA3G (**Figure** 3A-B). There, HIV-2 Vif encodes high backbone flexibility that expands the range of conformational sampling, thereby allowing HIV-2 Vif tryptophan sidechains to form more extensive interactions with hA3G than HIV-1 Vif (**Figure** 4). When the Trp-Rich motif of HIV-2 Vif is restrained by the preceding residue, HIV-2 Vif G48P, there is a modest reduction in infectivity levels against hA3G (**Figure** 5B). We propose a model where flexible loops in HIV-2 Vif undergo conformational selection to bury additional surface area and allow recognition of diverse A3Gs, whereas HIV-1 Vif binds a smaller interface on hA3G through a more rigid lock-and-key mechanism.

### SIVrcm Vif Trp-Rich Loop Contributes to Modest Neutralization of hA3G

Although HIV-2 and SIVrcm Vif have similar Trp-Rich loops, SIVrcm Vif is unable to robustly neutralize hA3G (**Figure** 3C, 3E, and 5A). Our study demonstrates the Trp-Rich loop in SIVrcm Vif has reduced backbone flexibility compared to HIV-2 Vif, likely due to conformational restraints imposed by the proline preceding the Trp-Rich motif (**Figure** 4A-B). When the Trp-Rich loop in SIVrcm Vif is further constrained by substitution of the glycine spacing the Trp-Rich motif to an alanine, there is an over 10-fold reduction in infectivity against hA3G (**Figure** 5A). These data suggest SIVrcm Vif may form a Trp-Rich interface with hA3G, but with weaker interactions than HIV-2 Vif due to reduced loop conformational sampling. Outside of HIV-1 and SIVs infecting chimpanzees and gorillas, Vifs often include an aromatic-rich motif in loop 3 with high variation in the residues preceding and spacing the motif and we speculate an aromatic-rich Vif loop 3 be an important feature for A3G recognition across species (**Figure** S10A). Vifs must adapt to the A3Gs present in their host species and, perhaps, A3Gs of different species require unique compositions and conformations of an aromatic-rich loop in Vif. However, even when SIVrcm Vif encodes the same Trp-Rich loop as HIV-2 Vif, it maintains modest infectivity against hA3G (**Figure** 5A). Therefore, adaption in SIVrcm Vif’s Trp-Rich loop alone would be insufficient for cross-species transmission into humans.

SIVrcm Vif might recognize more diverse A3Gs if it encoded the same loop 5 sequence, with the accompanying loop dynamics, as HIV-2 Vif (**Figure** S9A-B). However, this type of adaptation in SIVrcm Vif loop 5 would represent a significant barrier as it would require nucleotide substitutions across three or more codons (**Figure** S6A and C). Instead, SIVrcm Vif underwent an adaption in loop 5 (Y86H) which requires only a single-nucleotide polymorphism (TAT to CAT, or TAC to CAC) and allows near full antagonism against hA3G ^14^. After adaptation in Vif loop 5 occurred, Vifs in the HIV-1 lineage may have lost the Trp-Rich motif as there was no longer a functional advantage for maintaining the tryptophans. Currently, there are no structures of SIVrcm Vif bound to an A3G and our study suggests there may be other determinants for A3G antagonism beyond SIVrcm Vif loop 5. Further studies are needed to elucidate the role of SIVrcm Vif’s Trp-Rich loop, as well as loop flexibility, in A3G recognition.

### HIV-2 Vif’s Arms Race and Trp-Rich Loops Multitask for A3H Recognition

A recent structural study demonstrated HIV-2 Vif utilizes the same interface (loops 3 and 5) to recognize both hA3G and human APOBEC3H haplotype II (hA3H hapII), a haplotype that encodes a more stable antiviral protein ^20–23^. Upon mutation to its Trp-Rich motif (GWAWW to GASAW), HIV-2 Vif loses function against hA3H hapII but retains some activity against hA3G ^17,23^. While HIV-2 Vif loop 5 adopts similar a conformation in these structures, HIV-2 Vif Y85 points away from hA3G and towards hA3H hapII ^23^. Thus, structural plasticity of HIV-2 Vif loop 5 may be necessary for neutralization of both hA3G and hA3H hapII, which is in line with our model of conformational selection in HIV-2 Vif for A3 recognition (**Figure** S11).

While A3H has repeatedly lost its antiviral activity throughout primate evolution, including in many Old World monkey and human populations, active forms of A3H represent a significant barrier to cross-species transmissions (Garcia & Emerman, 2018; Zhang et al., 2017). As tryptophan is encoded by only one codon and can be easily lost through single-nucleotide polymorphism, the conservation of the GWAWW motif in SIVsmm Vif and its descendants (HIV-2 and SIVs from macaques) suggests that this motif is functionally important (**Figure** S10B). Notably, both sooty mangabey monkeys and rhesus macaques encode for active A3H ^24^. Since the same surface on HIV-2 Vif is used to recognize diverse A3s, the evolutionary pressure from active A3H alleles may explain the conservation of the GWAWW motif in the SIVsmm Vif lineage.

### Flexible Loops in Vif Can Encode Broad Specificity

As protein-protein interactions are often mediated through loop regions, it is critical to understand how loop dynamics tune specificity. The conformational ensembles of loops are often related to substrate recognition, with loop flexibility often being a trade-off between breadth and affinity. For example, the complementarity-determining region (CDR) loops of antibodies are often rigidified during affinity maturation to increase binding affinity and specificity, as flexibility comes at entropic costs as CDRs become ordered upon antigen binding ^25–27^. However, this entropic cost can be worthwhile in the pursuit of broadly neutralizing antibodies as flexible CDRs can accommodate mutated or otherwise diverse antigens ^28–30^. Similarly, HIV-2 Vif recognizes diverse A3s, A3G and A3H, with two loops exhibiting more dynamic backbones than the equivalent loops in HIV-1 (**Figure** 4). As RNA acts a molecular glue between Vif and hA3G, interactions at the RNA-mediated interface may compensate for the entropic cost of Vif loop flexibility when binding A3Gs with sequence variation ^6–8^. However, the Vif flexibility required for broad A3 recognition may result in poor binding affinity for other A3s and lower infectivity levels. Compared to the HIV-1 lineage, HIV-2 and its precursor virus arose more recently ^31^. As HIV-2 Vif and human A3s continue to coevolve, HIV-2 Vif loops may become rigidified over time in favor of higher binding affinities for the A3s that represent the most significant barriers to viral transmission, and broad A3 recognition may be lost.

## Methods

### Cell Lines

Sf9 and Hi5 insect cells were maintained in Sf-900 III SFM medium (Gibco) at 27°C with shaking at 120 rpm. HEK293T/17 cells were maintained in Dulbecco’s modified Eagle’s medium (DMEM, Thermo Fisher Scientific) supplemented with 10% fetal bovine serum (FBS, Corning) and 1% Antibiotic-Antimycotic (Gibco) at 37°C in a humidified incubator with 5% CO_2_. SupT1 cells were cultured in RPMI 1640 medium (Gibco) containing 10% FBS, 1% Antibiotic-Antimycotic, 1% sodium pyruvate, 1% glucose, and 1% GlutaMAX (Gibco)under the same conditions.

### HIV-2 Vif Protein Expression Screen and Western Blot

Six HIV-2 Vif sequences, five of which were from patient isolated strains (accession numbers: AF082339, M31113, U27200, KY025545, and L07625) and one of which was from a laboratory adapted strain (HIV-2_ROD_, accession number: M15390), were selected for expression tests in insect (Sf9) cells. The HIV-2 Vif sequences were codon optimized and fused to a C-terminal FLAG tag. Then, each HIV-2 Vif sequence was cloned into a single MacroBac vector with human A3G fused to a C-terminal 2xStrep tag, CBFβ, EloB, EloC, and CUL5 residues 1-386 (referred to as CUL5N) as previously described ^32^. These six constructs were screened for co-expression in Sf9 insect cells. Baculoviruses were generated with the Bac-to-Bac system and amplified in Sf9 cells. Small-scale protein expression (25mL) was performed in Sf9 cells. Cells were harvested 48 h post-infection in 6 mL aliquots by centrifugation at 2,500 x g for 10 min at 4°C at -80°C until use. Each 6 mL cell pellet was resuspended and lysed in 100uL of 1% (v/v) Triton X-100 and re-pelleted at 17,000xg for 10 min at room temperature to remove cell debris. The clarified cell lysate samples were diluted with 4X loading dye and boiled for ten minutes at 95°C before being loaded onto a 4-15% SDS-PAGE gel then transferred onto a nitrocellulose membrane with the Trans-Blot Turbo system (Bio-Rad). Anti-Strep HRP-conjugated (Sigma-Aldrich) and rabbit anti-FLAG (Sigma-Aldrich) antibodies were used for immunoblotting at respective dilutions of 1:5,000 and 1:2,500. Afterward, StarBright Blue 520 Goat Anti-rabbit IgG (Bio-Rad) was used at a dilution of 1:5,000. Fluorescent and chemiluminescent signals were detected with a ChemiDoc MP Imaging System (Bio-Rad).

### Protein Expression and Purification

HIV-2_ALI_ Vif (Accession number: AF082339) was selected for further study as it expressed well in Sf9 cells and exhibits strong neutralization of hA3G that is comparable to HIV-2_ROD_ Vif. Large-scale protein expression was performed in Hi5 cells infected with baculovirus. Cells were harvested 48 h post-infection by centrifugation at 2,500 x g for 20 min at 4°C. Cell pellets were washed once with PBS, flash-frozen in liquid nitrogen and stored at -80 °C until use. Cell pellets from 1 L of Hi5 culture were resuspended in 50 mL lysis buffer (50 mM HEPES pH 8.0, 50 mM NaCl, 5% (v/v) glycerol, 1% (v/v) Triton X-100, 5 mM MgCl_2_, 5 mM CaCl_2_, 1 mM TCEP, cOmplete protease inhibitor tablets (5 tablets, Sigma-Aldrich), 25 µg mL^-1^ DNase I (Sigma-Aldrich), and 50 µg mL^-1^ RNase A (Sigma-Aldrich)). Cells were lysed using a Dounce homogenizer and incubated on ice for 30 min before centrifugation at 17,000 x g for 1.5 h at 4°C to remove debris. The clarified lysate was passed through a 0.45 µm filter and batch incubated for 1 h at 4°C with 5 mL Strep-Tactin Sepharose resin (IBA Lifesciences) pre-equilibrated with binding buffer (50 mM HEPES pH 8.0, 150 mM NaCl, 5% (v/v) glycerol, and 1 mM TCEP). The resin was washed three times with 5 CVs of wash buffer (50 mM HEPES pH 8.0, 500 mM NaCl, 5% (v/v) glycerol, and 1 mM TCEP) and one time with 5 CVs of binding buffer. The complex was eluted with 6 CVs of elution buffer (50 mM HEPES pH 8.0, 150 mM NaCl, 5% (v/v) glycerol, 1 mM TCEP, and 50 mM D-desthiobiotin). The eluate was concentrated and dialyzed in 1 L of dialysis buffer (50 mM HEPES pH 7.0, 150 mM NaCl, 5% (v/v) glycerol, and 1 mM TCEP) overnight at 4°C. After dialysis, the complex was diluted two-fold with salt free dialysis buffer (50 mM HEPES pH 7.0, 5% (v/v) glycerol, and 1 mM TCEP) and loaded onto a 5 mL HiTrap Heparin HP column (GE Healthcare) pre-equilibrated with buffer A (50 mM HEPES pH 7.0, 75 mM NaCl, 5% (v/v) glycerol, and 1 mM TCEP). The complex was eluted using a linear NaCl gradient from 75 mM to 1 M. Fractions containing all six components were pooled and further purified by size-exclusion chromatography on a Superose 6 Increase 10/300 GL column (GE Healthcare) pre-equilibrated with sizing buffer (30 mM HEPES pH 7.0, 150 mM NaCl, 5% (v/v) glycerol, and 1 mM TCEP). The peak fraction was used for cryo-EM sample preparation and HDX-MS.

SIVrcm Vif, rcmA3G fused to a C-terminal 2xStrep tag, CBFβ, EloB, EloC, and CUL5N were cloned into a single MacroBac vector, co-expressed in insect cells, cell pellets were harvested and lysed as described above. 1 mg of NeutrAvidin (Thermo Fisher Scientific) was added to the cell lysate before centrifugation at 17,000 x g for 1.5 h at 4°C to remove debris. The clarified lysate was passed through a 0.45 µm filter and batch incubated for 2 h at 4°C with 4 mL Strep-Tactin Sepharose resin (IBA Lifesciences) pre-equilibrated with binding buffer. The resin was washed with 50mL of wash buffer, followed by 50mL of binding buffer. Then the resin was incubated overnight at 4°C with 20mL of binding buffer supplemented with 50 µg mL^-1^ RNase A. The resin was washed with another 50mL of wash buffer and the complex was eluted with 30mL of elution buffer. The elution was diluted with 50mL of buffer A and then loaded onto a 5 mL HiTrap Heparin HP column (GE Healthcare) pre-equilibrated with buffer A. The complex was eluted using a linear NaCl gradient from 75 mM to 1 M. Fractions containing all six components were pooled and further purified by size-exclusion chromatography on a Superose 6 Increase 10/300 GL column (GE Healthcare) pre-equilibrated with sizing buffer. The peak fraction was used for HDX-MS.

HIV-1_HXB2_ Vif, hA3G with a C-terminal 2xStrep tag, CBFβ, and EloB, EloC, and CUL5N were co-expressed and purified as previously described (Y. L. Li et al., 2023). The peak fraction was used for HDX-MS.

### Cryo-EM Grid Preparation and Data Collection

The purified hA3G, HIV-2_ALI_ VCBC, and Cul5N complex was used directly for cryo-EM sample preparation, with an A_280_ of 0.314 mg mL^-1^ corresponding to an estimated concentration of 2 µM. Aliquots of 3.5 µL were applied to glow-discharged Quantifoil R 1.3/1.3 300 Mesh Au grids (Electron Microscopy Sciences). Grids were blotted with a wait time of 15 s, blot time of 12 s, and blot force of 0 at 23°C and 100% humidity using a Vitrobot Mark IV (Thermo Fisher Scientific), then plunge-frozen in liquid ethane that was cooled by liquid nitrogen.

Cryo-EM grids were screened on a 200 kV Arctica microscope (Thermo Fisher Scientific). High-quality grids were selected for data collection on a 300 kV Titan Krios microscope (Thermo Fisher Scientific). 17,753 movies were recorded on a Gatan K3 direct electron detector mounted post a quantum energy filter (20 eV slit width) at a magnification of 105,000x and pixel size of 0.4094 Å. Data were collected using SerialEM with defocus values ranging from -1 µm to -2.2 µm, a dose rate of 16 e^−^/pixel/s, and an exposure time of 2 s (0.025 s per frame), yielding 80 frames per stack and a total dose of 47.7 e^−^/Å^2 34^. Movie stacks were motion-corrected using MotionCor2 within the SCIPION framework and binned two-fold, resulting in a pixel size of 0.8189 Å ^35,36^.

### Cryo-EM Image Processing and Model Building

Dose-weighted, motion-corrected micrographs were manually curated, where ones with excessive ice contamination or poor hole targeting were excluded, and then imported into cryoSPARC 4.4.1 for data processing ^37^. The contrast transfer function (CTF) of the micrographs was estimated by Patch CTF and the dataset was further manually curated to 17,404 micrographs. Using reference-free Blob Picker (minimum particle diameter of 150Å, maximum particle diameter of 200Å), 9,552,691 particles were automatically picked and 5,387,868 particles were selected for particle extraction. Particles were extracted in a 352-pixel box size and binned four-fold to 88-pixels (3.2756Å per pixel). Two rounds of 2D classification were performed and classes with proteinaceous features were selected for further processing. These 2,258,305 particles were subjected to heterogeneous refinement with three reference maps (one generated from PDB 8CX0 and low pass filtered to 20Å and two junk). The 798,711 particles from the best class (reference: map generated from PDB 8CX0) were re-extracted from the micrographs without binning. These re-extracted particles were subjected to further heterogeneous refinement with four reference maps (generated from PDB 8CX0, 8CX1, and 8CX2 each low pass filtered to 20Å and one junk). The class of the monomeric complex (reference: map generated from PDB 8CX0) containing 357,465 particles was used for non-uniform refinement (to generate a reference for further refinement) and one round of additional 2D classification where only classes with strong proteinaceous features were selected. These 152,430 particles were refined with non-uniform refinement before being re-extracted for migration to the RELION software ^38^. These computational analyses were performed using the SBGrid software environment, which provides curated, version-controlled structural biology applications and reproducible execution environments across platforms ^39^. This particle stack was further refined using Refine3D with Blush Regularization and CTF Refine. To further improve the resolution, a mask was generated and applied during another round of Refine3D with Blush Regularization and the map was sharpened in RELION-5.0. 3DFSC was used to calculate the global resolution of the final masked map (3.6Å) and local resolution this map was calculated in RELION-5.0 ^38,40^.

The AlphaFold2 predicted structure of HIV-2_ALI_ VCBC and the cryo-EM structure of hA3G bound to RNA (PDB 8CX0) were docked into the map with ChimeraX-1.11.1 for initial model building of the complex ^6,41–43^. The model went through iterative rounds of manual adjustments in Coot-0.9.8.6 and refinement with real space refine in PHENIX-1.19.2 with secondary structure and geometry restraints applied ^44,45^. The final model was validated using the cryo-EM validation tool in Phenix ^45^. All structure figures were prepared using ChimeraX-1.11.1 ^43^.

### Viral Infectivity Assay

Single cycle infectivity assays were performed as previously described ^23^. Untagged WT or mutant Vif sequences were cloned into the pLAIΔenvLuc2ΔVif proviral vector. HEK293T/17 cells were seeded at 1.5 x 10^5^ cells/mL in 6-well plates and transfected with 400 ng of the proviral vector, 100 ng of L-VSV-G and 200 ng of an N-terminal HA-tagged A3G (WT hA3G, D128K hA3G, or rcmA3G) or empty vector control plasmid using TransIT-LT1 transfection reagent (Mirus Bio) at a ratio of 3.5µL reagent per µg DNA. At 24h post-transfection, media were replaced. Viral supernatants were harvested at 72h post-transfection and clarified by filtration through 0.22-µm PES filters. Viral titers were quantified by RT-qPCR based titration ^46^. Equal infectious units were used for all infections, with mock-transfected supernatant added to equalize final volumes.

SupT1 cells were seeded at 1.5 x 10^5^ cells/mL in U-bottom 96-well plates in media supplemented with 20 µg/mL DEAE-dextran and infected by spinoculation at 1100 x g for 30 min at 30 °C. All infections were performed in technical duplicates, and each condition was tested in three independent biological replicates. At 72 h post-infection, cells were lysed with 100 µL Bright-Glo reagent (Promega), and luminescence was measured using a LUMIstar Omega luminometer (BMG Labtech). The percent infectivity was calculated by normalizing the relative light units (RLUs) from each biological replicate containing A3G protein to the mean RLUs of the corresponding virus produced in the absence of A3G.

### Quantification and Statistical Analysis

The single-cycle infectivity data are presented as mean ± SD from at least three independent replicates. Statistical analyses were performed using GraphPad Prism 11, and significance was determined using unpaired Welch’s t-tests (*p < 0.05, **p < 0.01, ***p < 0.001, ****p < 0.0001).

### Hydrogen-deuterium exchange

Biochemical and statistical details for all HDX experiments are summarized in Supplementary Table 2. HDX was performed on three complexes: (1) HIV-1 VCBC/hA3G, (2) HIV-2 VCBC/hA3G, and (3) SIVrcm VCBC/rcmA3G. Labeling was initiated by diluting 10 μL of 2 μM complex in H_2_O buffer (30 mM HEPES, 150 mM NaCl, 5% glycerol, 1 mM TCEP, pH 7.0) into 30 μL of matching deuterated buffer (71% D-content final) at pD_read_ 6.6 and temperatures of either 2°C or 25°C. Labeling times were adjusted to account for the temperature dependence of intrinsic rates (*k*_*chem*_), calculated from the average *k*_*chem*_ of the full sequence ^47^. Selected timepoints were collected in triplicate.

HDX was quenched at time points ranging from 3 sec to 24 hours by adding 40 μL of ice-cold quench buffer (600 mM Glycine, 4 M urea, pH 2.5) and then immediately injected into a valve system at 2°C (Trajan LEAP). Non-deuterated controls and MS/MS runs for peptide assignment followed the same protocol with H_2_O substituted for D_2_O and immediate quenching and injection. HDX reactions were performed in random order. No peptide carryover was observed as assessed by injecting quench buffer containing 2 M urea. Maximally labeled controls accounting for back-exchange were prepared by incubating samples in a D_2_O buffer at pD_read_ 13 for 1 h at 22°C.

### Liquid chromatography – mass spectrometry

Protease columns were selected to maximize sequence coverage and redundancy at Loop 3 and Loop 5. HIV-1 VCBC/hA3G and HIV-2 VCBC/hA3G complexes were digested online with a NepII/pepsin protease column (Affipro AP-PC-006) at 10°C; SIVrcm VCBC/rcmA3G was digested with a pepsin/FPXIII mixed protease column at 10°C, prepared in-house by coupling the protease (Sigma-Aldrich P6887/P2143) to a resin (Thermo Scientific POROS 20 Al aldehyde activated resin 1602906) and hand packing into a column (2 mm ID × 2 cm, IDEX C-130B). The resulting peptides were desalted on a C8 trap column (Waters ACQUITY UPLC BEH C8 VanGuard Pre-column, 130Å, 1.7 µm, 2.1 mm X 5 mm, 186003978) at 2 °C. Digestion and desalting were completed in 3 min at 150 μL/min of 0.1% formic acid. Peptides were then separated on a C18 analytical column (Waters ACQUITY UPLC BEH C18, 130Å, 1.7 µm, 1 mm X 50 mm,186002344) using a 17-min linear gradient of 5–45% (vol/vol) acetonitrile (0.1% formic acid) delivered by a Vanquish Neo pump. Eluted peptides were analyzed on a Thermo Q Exactive Plus mass spectrometer in positive mode with following settings. Peptide assignment: full MS: resolution 70,000, AGC target 3 x 10^6^, max IT 100 ms, scan range 300-1500 m/z; dd-MS^2^: resolution 35,000, AGC target 1 x 10^5^, max IT 100 ms, loop count 10, isolation window 2.0 m/z, NCE 28, dynamic exclusion 15 sec. For HDX: full MS: resolution 140,000, AGC target 3 x 10^6^, max IT 200 ms, scan range 300-1500 m/z.

### HDX-MS data analysis

Peptides were identified using SearchGUI (v4.0.25) against a library containing HIV-1 Vif, HIV-2 Vif, SIVrcm Vif, hA3G, rcmA3G, CBFβ, EloB, and EloC, plus 15 additional proteins previously exposed to the LC-MS system, with reserved sequences as decoys. Search parameters included unspecific digestion, precursor and fragment tolerance 10 ppm, precursor charge 1-8, and peptide length 5-50 residues. HDX data were processed in HDExaminer 3.5.0 (Trajan). Deuteration levels were corrected for 71% D-content and back-exchange.

HDX in this study occurred via EX2 kinetics, where the observed exchange rate reports on the equilibrium (i.e., stability) rather than the opening rates of the exchange-competent states. EX2 behavior was confirmed by continuous shifts in unimodal isotopic envelopes toward higher m/z with exchange time.

Residue-level unfolding free energies (ΔG) for Loop 3 and Loop 5 were calculated as follows. Non-overlapping atomic ranges were determined by HDExaminer from all overlapping peptides, and per-range deuteration occupancy were computed by minimizing least-squares error against the measured peptide deuteration levels. All isotope distribution and deuteration fits were manually inspected for fit quality. Deuteration levels for each residue were then fit to:

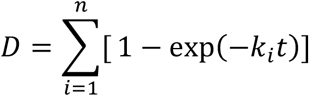

where *k*_*i*_ is the exchange rate for each residue, and *n* is the number of residues in the atomic range. For atomic ranges containing more than one residue, a single exchange rate was calculated by averaging the individual exchange rates within that range. Under EX2 limit, the unfolding free energy (ΔG) was calculated as

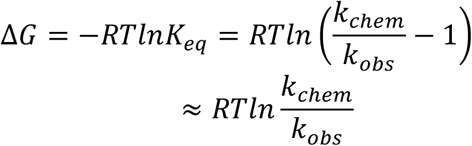

Residue-level unfolding free energies (ΔG) for hA3G were calculated using a stretched exponential method described in ^48^.

### Vif Sequence Analysis

Sequences of natural variants of SIV, HIV-1, and HIV-2 Vif were obtained from premade alignments curated by the Los Alamos HIV database (https://www.hiv.lanl.gov/). Logo plots were generated using WebLogo 3 ^49^. The number of unique sequences per SIV or HIV lineage are referenced in the figure legends of each corresponding logo plot.

## Data Availability

The cryo-EM map and atomic model have been deposited in the EMDB and PDB under accession codes EMD-77600 and PDB 36IM, respectively. Raw cryo-EM data (motion corrected micrographs, particle stack, unaligned multi-frame micrographs, and gain reference) have been deposited to the EMPIAR database under EMPIAR-13950. The HDX mass spectrometry data have been deposited to the ProteomeXchange Consortium via the PRIDE ^50^ partner repository with the dataset identifier PXD083821. Further information and requests for resources and reagents should be directed to and will be fulfilled by the corresponding authors, Dr. John D. Gross and Dr. Yifan Cheng.

## Acknowledgements

We thank members of the Gross, Cheng, and Emerman laboratories for invaluable discussions and sharing their expertise, especially Yen-Li Li, Kevin Choi, and Amber Smith. We are grateful to David Bulkley, Glenn Gilbert, and Li Wang at the UCSF Cryo-EM facility for their assistance with cryo-EM data acquisition.

## Funding

This research was supported by National Institutes of Health (NIH) and National Institute of Allergy and Infectious Diseases (NIAID) funding for the HIV Accessory and Regulatory Complexes (HARC) Center (U54AI170792 to J.D.G., M.E., and Y.C.), as well as additional NIH funding to J.D.G. and N.M.C. (R01AI202673) and Y.C. (R35GM140847). E.K.R. was supported by the National Science Foundation (NSF) Graduate Research Fellowship Program (Grant No. 2445150) and University of California, San Francisco’s Discovery Fellowship. Y.N. was supported by a Mentored Scientist Award from the UCSF-Bay Area Center for AIDS Research (CFAR), funded by the National Institutes for Health (P30 AI027763), and a Mentored Scientist Award from the HARC Center (U54AI170792). The UCSF Cryo-EM facility and its instruments are supported partially by grants from the NIH (S10OD020054, S10OD021741, and S10OD026881) and Howard Hughes Medical Institute (HHMI). Y.C. is an HHMI Investigator.

## Author Contributions

E.K.R., Y.C. and J.D.G conceived the project. Y.C. and J.D.G. supervised the research. E.K.R. expressed and purified the HIV-1 and HIV-2 protein complexes, collected and processed cryo-EM data, and performed model building and refinement. M.L. performed infectivity assays and analyzed the data under the supervision of N.M.C. and M.E. X.L. performed HDX-MS and processed the data. Y.N. expressed and purified the SIVrcm protein complex. E.K.R., X.L., and M.L. created the figures. All authors contributed to data interpretation. E.K.R. wrote the manuscript with input from all authors.

## Competing interests

Y.C. is a non-shareholder member of the scientific advisory boards for ShuiMu BioSciences and Pamplona Therapeutic Co. The other authors declare no competing interests.

## Correspondence and requests for materials

should be addressed to Dr. Yifan Cheng and Dr. John D. Gross.

---

**Figure S1.**
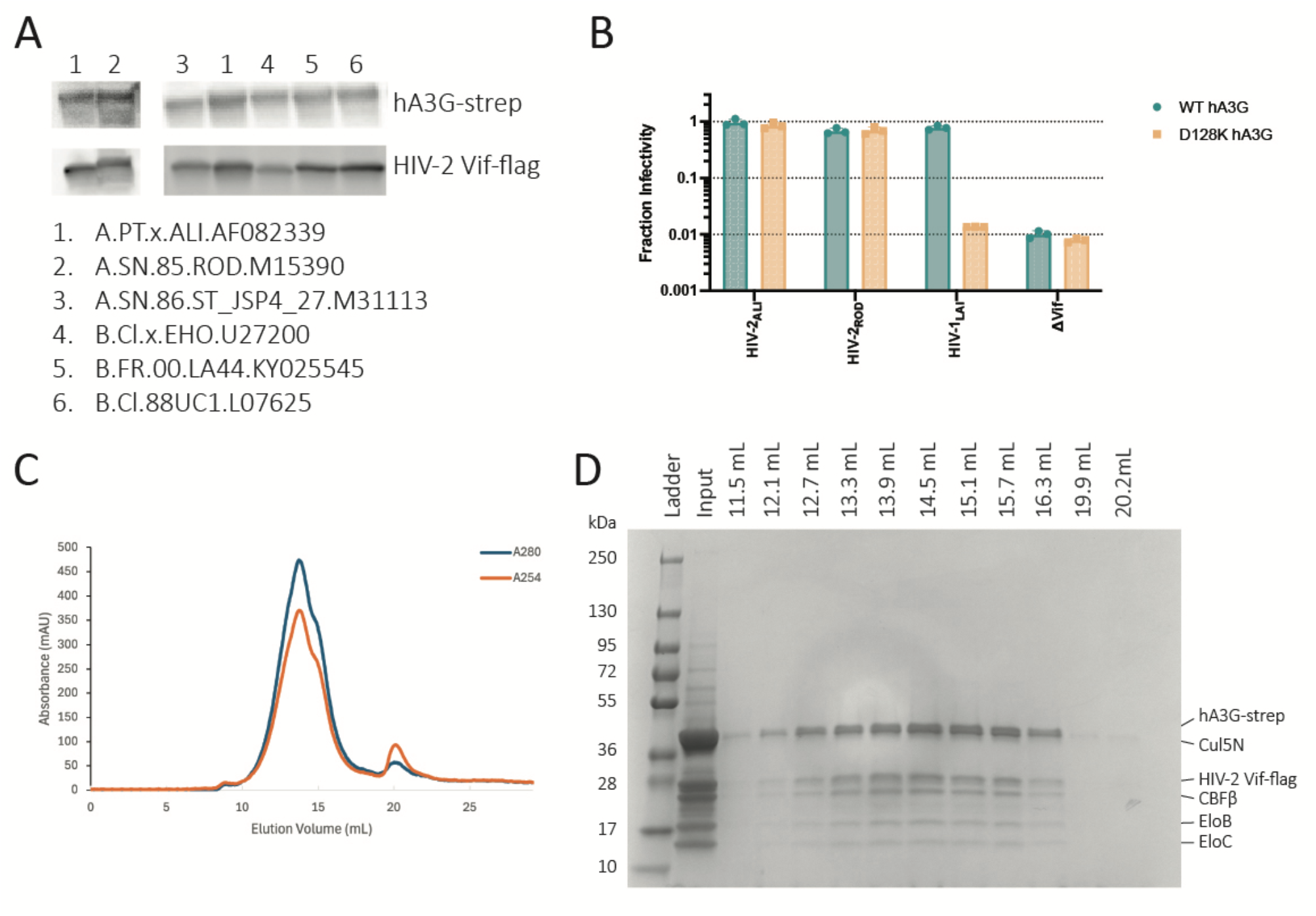
Screen of HIV-2 Vif strains and purification of the hA3G and HIV-2 VCBC complex. **A)** Western blot comparing hA3G-strep and HIV-2 Vif-flag expression for five patient isolated HIV-2 Vif sequences and laboratory adapted HIV-2_ROD_ Vif. Vif sequences were screened for co-expression with hA3G, CBFβ, EloB, EloC, and Cul5N in the baculovirus-insect cell expression system. **B)** Single-cycle viral infectivity assay of HIV-2_ALI_ Vif, HIV-2_ROD_ Vif, HIV-1_LAI_ Vif, and dVif in the presence of WT hA3G (teal) or D128K hA3G (yellow). The Y-axis represents the fraction infectivity. Standard deviation from the mean of three biological replicates is represented by error bars. **C)** Size-exclusion chromatography of the purified hA3G and HIV-2 V_ALI_CBC complex (hereafter HIV-2 VCBC). Absorbance at A_280_ and A_254_ is shown in blue and orange, respectively. **D)** SDS-PAGE of the purified hA3G and HIV-2 VCBC complex.

**Figure S2.**
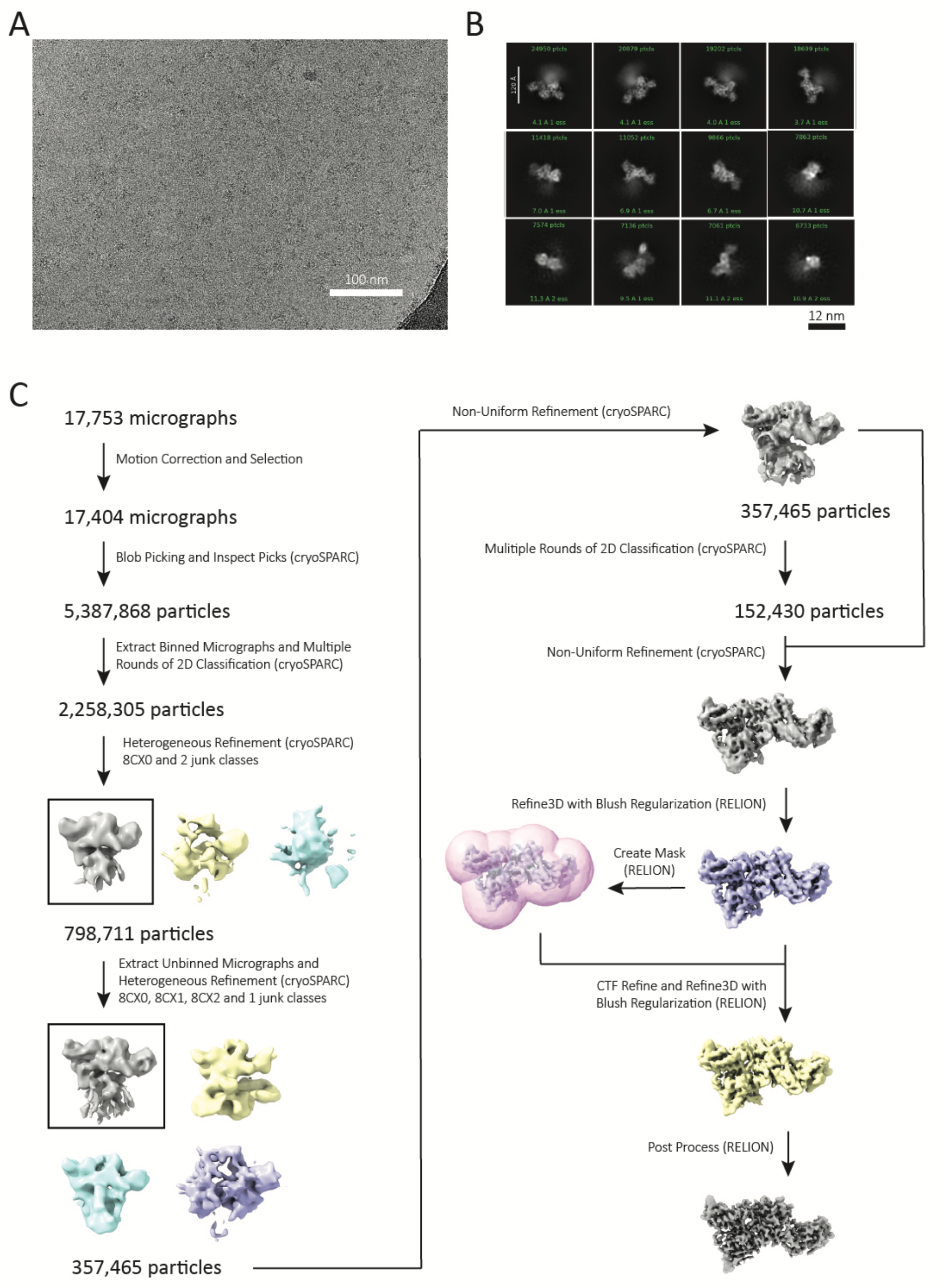
Cryo-EM data processing of the hA3G and HIV-2 VCBC complex. **A)** Representative cryo-EM micrograph. **B)** 2D class averages generated in cryoSPARC. **C)** Data processing workflow in cryoSPARC and RELION for reconstruction of the hA3G and HIV-2 VCBC complex.

**Figure S3.**
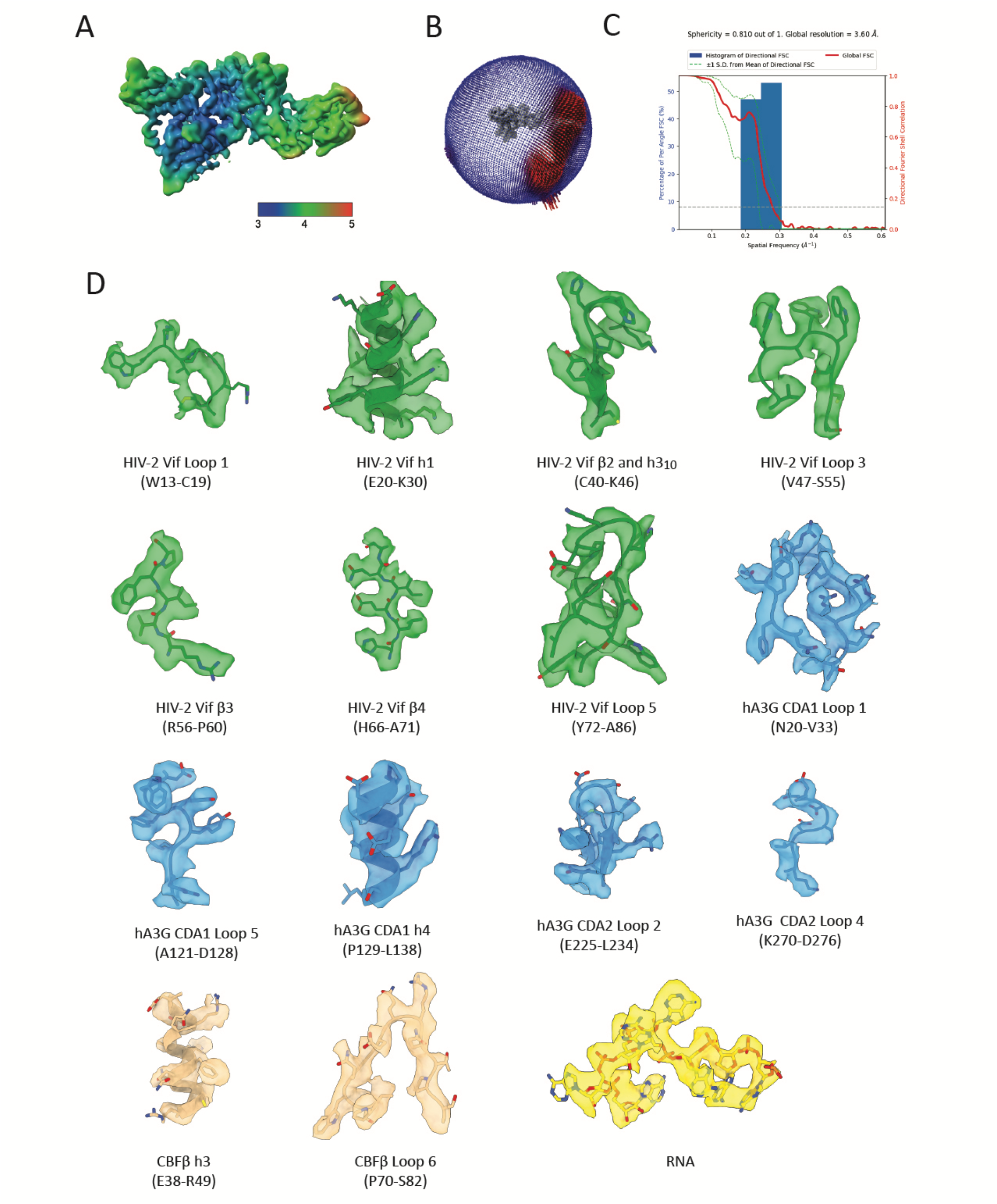
Cryo-EM quality metrics for the hA3G and HIV-2 VCBC complex. **A)** Local resolution calculated in RELION shown on the unsharpened map of the hA3G and HIV-2 VCBC complex. **B)** Angular distribution of particles. **C)** FSC Curve generated by 3DFSC. **D**. Sharpened cryo-EM map and refined atomic model for regions of interest. HIV-2 Vif, hA3G, CBFβ, and RNA are colored as in Figure 1.

**Figure S4.**
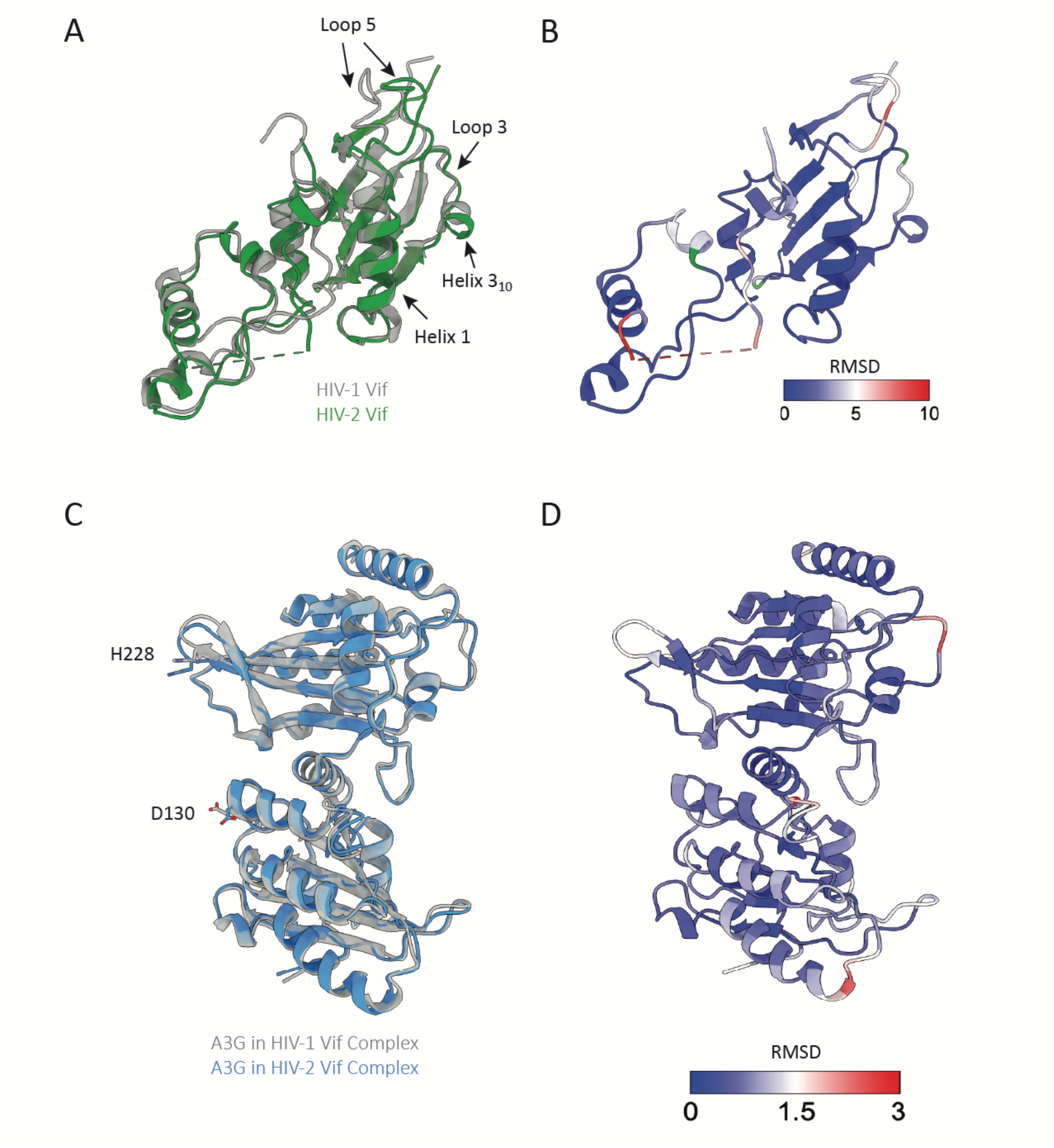
Structural comparison of HIV-2 Vif and hA3G with the HIV-1 complex. **A)** Alignment of HIV-2 Vif (green) and HIV-1 Vif (grey; PDB: 8CX0), with arrows pointing to regions of interest. **B)** RMSD between HIV-1 (PDB: 8CX0) and HIV-2 Vif plotted on the HIV-2 Vif structure. **C)** Alignment of hA3G from the HIV-2 complex (blue) and hA3G from the HIV-1 complex (grey; PDB: 8CX0), with D130 and H228 displayed as sticks. **D)** RMSD between hA3G in the HIV-1 (PDB: 8CX0) and HIV-2 complexes plotted on the hA3G structure from the HIV-2 Vif complex.

**Figure S5.**
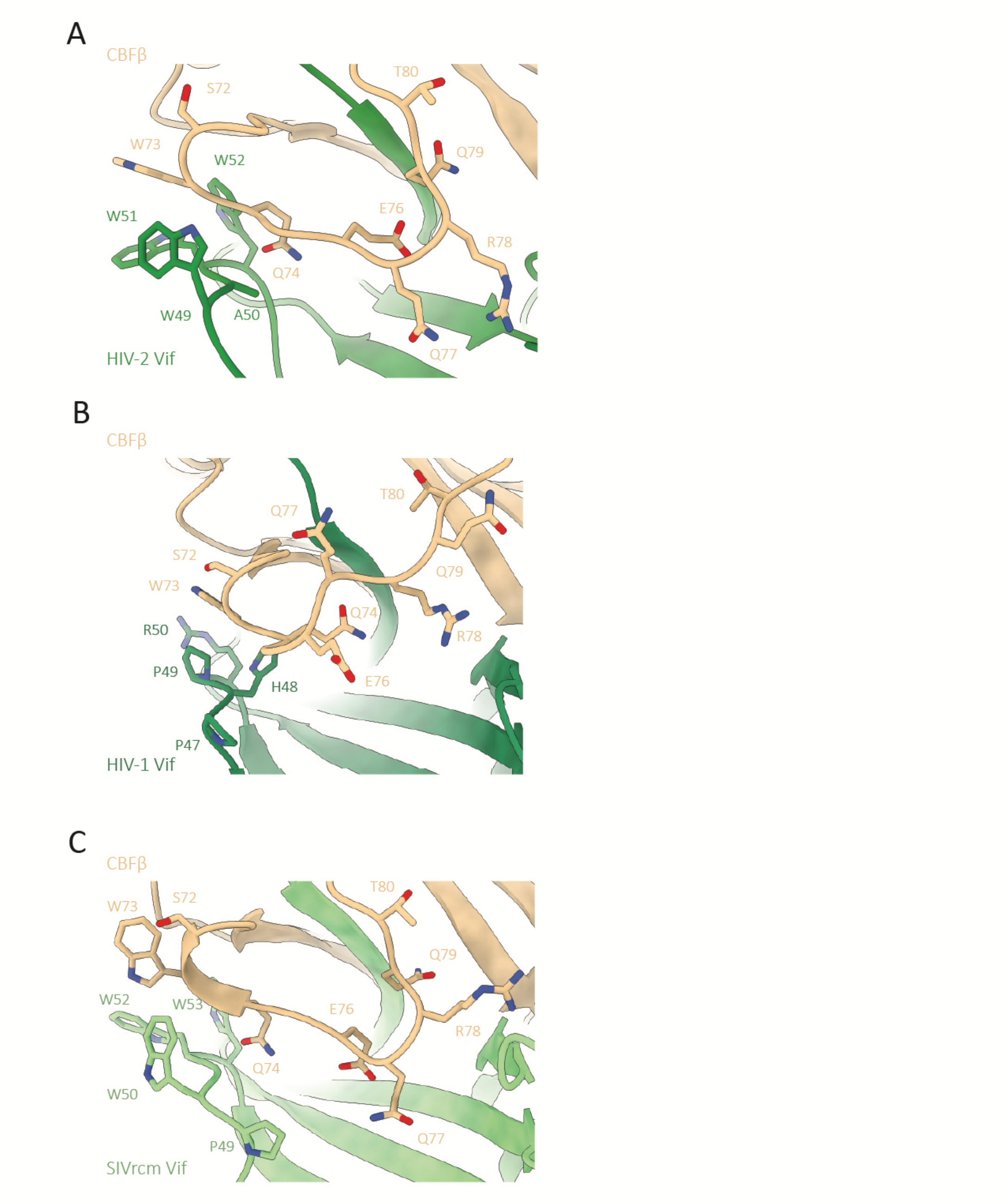
Structural comparison of CBFβ loop 6 in complex with HIV-2, HIV-1, or SIVrcm Vif. **A-C)** Closeup of where CBFβ (beige) loop 6 contacts Vif, with residues in Vif loop 3 and CBFβ loop 6 shown as sticks, for **A)** HIV-2 Vif (green). **B)** HIV-1 Vif (PDB: 8CX0; dark green) and **C)** SIVrcm Vif (PDB: 6P59; light green).

**Figure S6.**
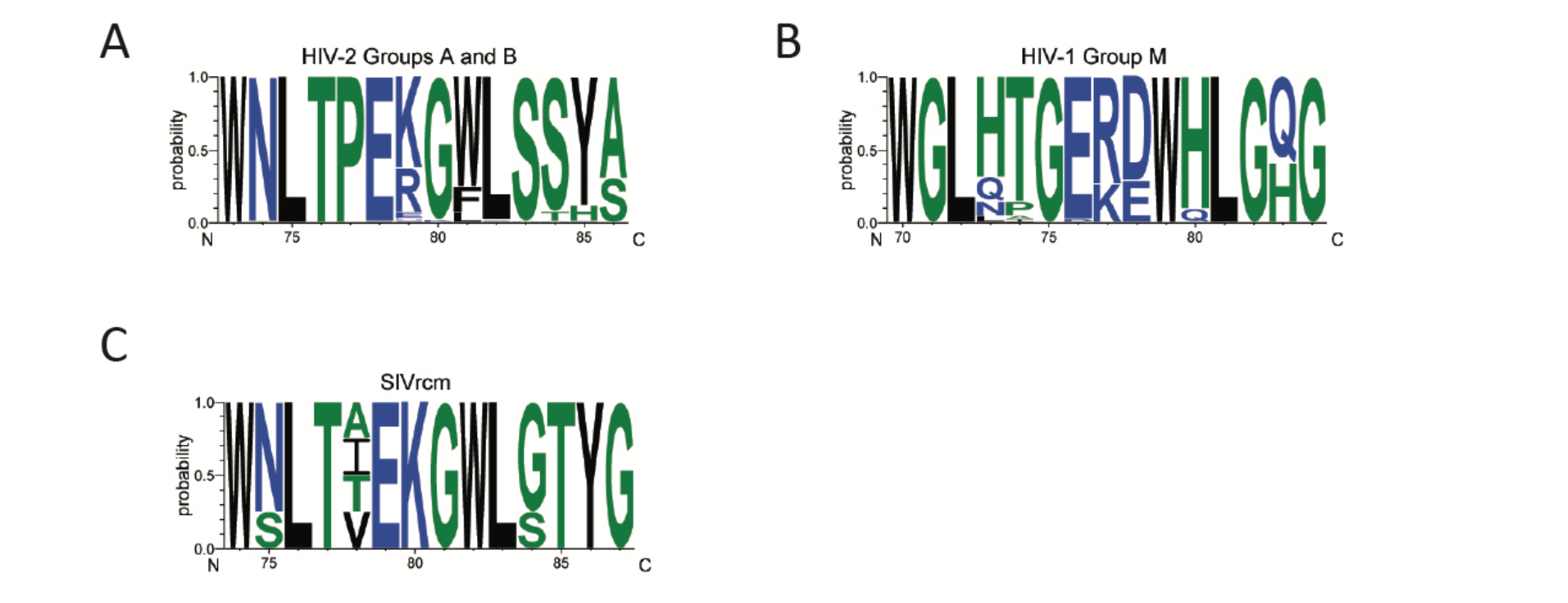
Sequence conservation of HIV-2, HIV-1, and SIVrcm Vif loop 5. **A-C)** Logo plot of sequence variation of Vif loop 5 for **A)** HIV-2 Groups A and B (104 sequences), **B)** HIV-1 Group M (42 sequences) and **C)** SIVrcm (4 sequences).

**Figure S7.**
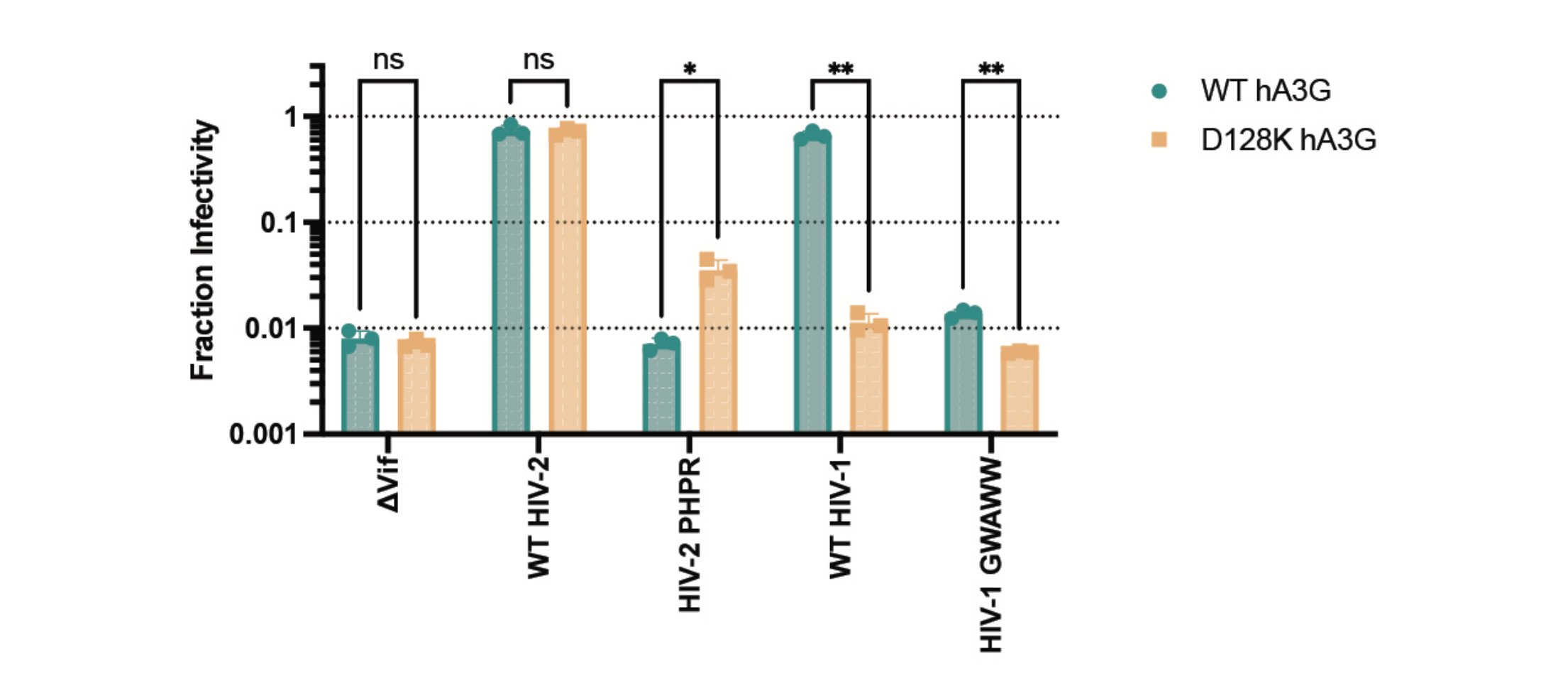
HIV-1 and HIV-2 loop 3 swaps result in loss of function. Single-cycle viral infectivity assay to assess the effect of HIV-1 and HIV-2 Vif loop 3 swaps in the presence of WT hA3G (teal) or D128K hA3G (yellow). The Y-axis represents the fraction infectivity. Standard deviation from the mean of three biological replicates is represented by error bars. * P < 0.05, ** P < 0.01, *** P < 0.001, **** P < 0.0001 as calculated by the unpaired Welch’s t-test.

**Figure S8.**
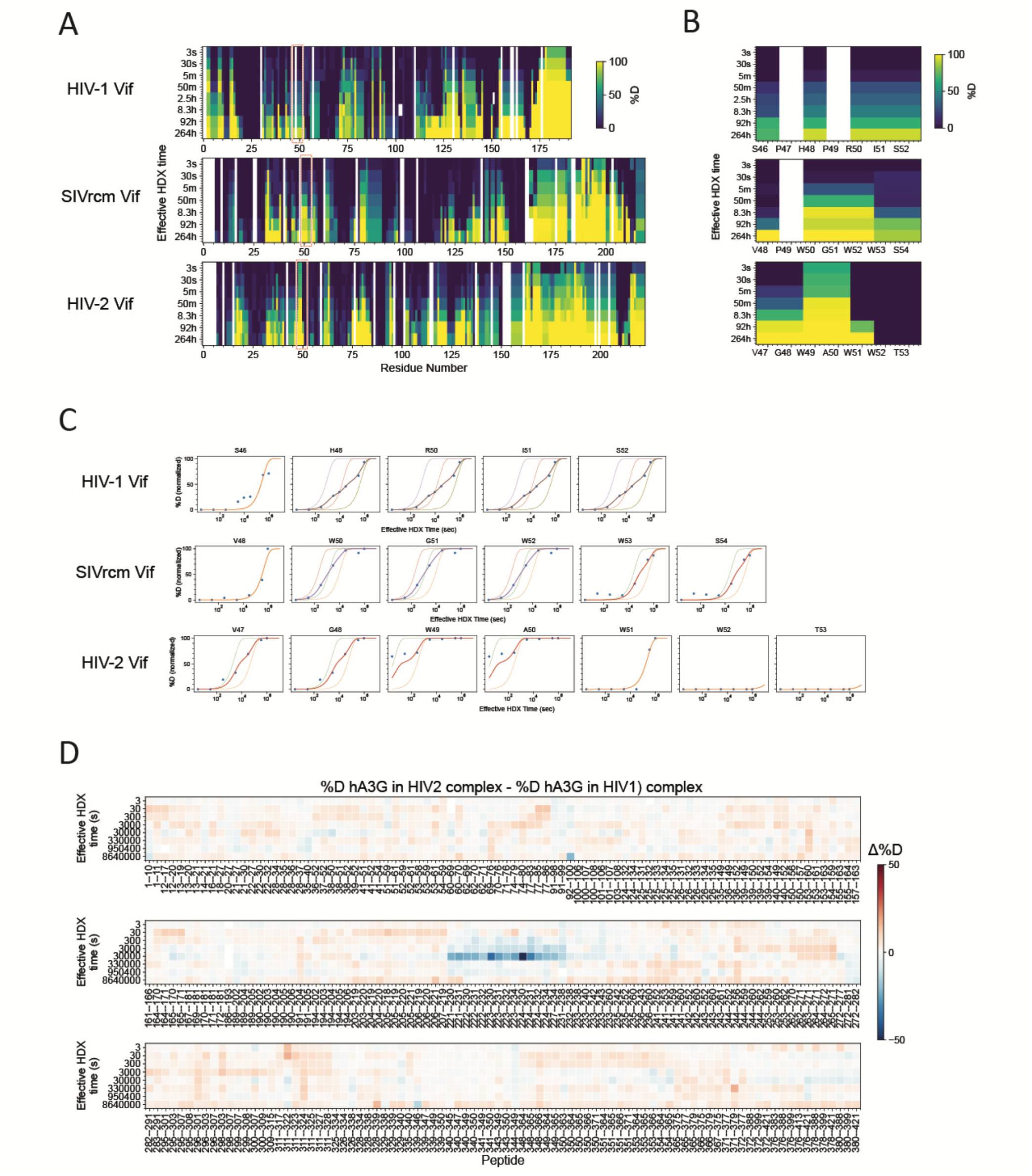
Stability of HIV-1 Vif, SIVrcm-Vif, HIV-2 Vif, and hA3G measured by HDX-MS. **A-B)** Heatmaps showing deuteration occupancy at each labeling time, computed for the smallest atomic ranges (HDExaminer). Red boxes in **A** highlight Loop 3, with enlarged views in **B. C)** Fitting results of Loop 3 deuteration kinetics. Exchange rates were determined by fitting a sum of exponentials to the labeling time courses. Solid circles: experimental data. Dashed lines: single exponential fit for individual residues when an atomic range spans more than one residue. Solid lines: sum-of-exponentials fit for the atomic range. **D)** Heatmaps showing changes in deuteration levels at each labelling time across all hA3G peptides, comparing the hA3G-HIV-2 Vif complex to the hA3G-HIV-1 Vif complex. Deuteration levels were corrected for 71% D-content and back exchange.

**Figure S9.**
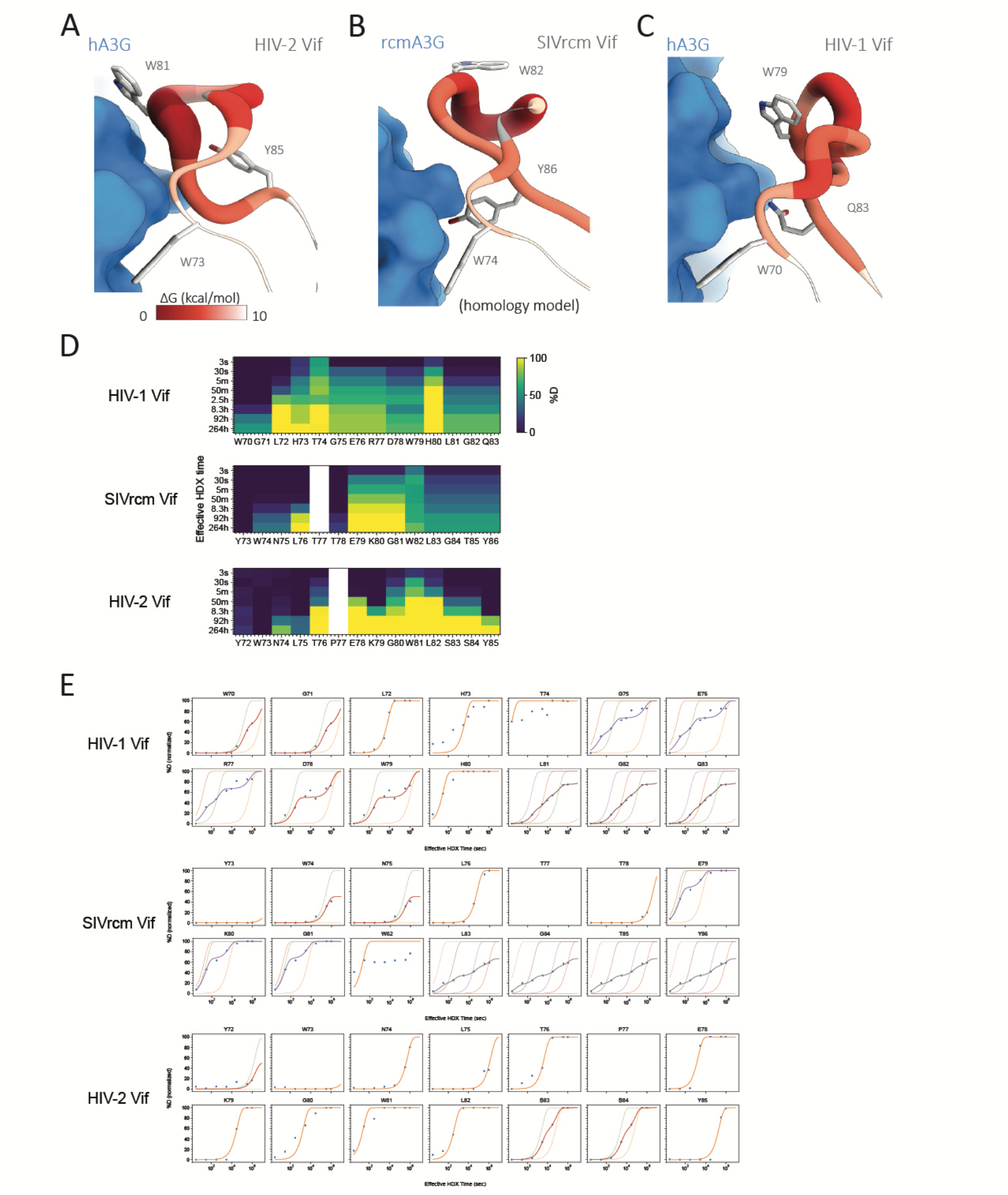
HIV-2 Vif encodes residues with low backbone stability at the arms race interface to engage hA3G. **A-C)** The Gibbs free energy of unfolding (ΔG) for **A)** HIV-2 Vif loop 5 while bound to hA3G, **B)** SIVrcm Vif loop 5 while bound to rcmA3G (visualized on a homology model of the rcmA3G and SIVrcm VCBC complex), **C)** HIV-1 Vif loop 5 while bound to hA3G (PDBID: 8CX0). A3G is shown as blue surface. Vif loop 5 is shown with backbone color and thickness reflecting ΔG values: red and thick for ΔG of 0 kcal/mol, grading to white and thin for ΔG of 10 kcal/mol. **D)** Heatmaps showing deuteration occupancy of loop 5 at each labeling time, computed for the smallest atomic ranges (HDExaminer). **E)** Fitting results of Loop 5 deuteration kinetics. Exchange rates were determined by fitting a sum of exponentials to the labeling time courses. Solid circles: experimental data. Dashed lines: single exponential fit for individual residues when an atomic range spans more than one residue. Solid lines: sum-of-exponentials fit for the atomic range.

**Figure S10.**
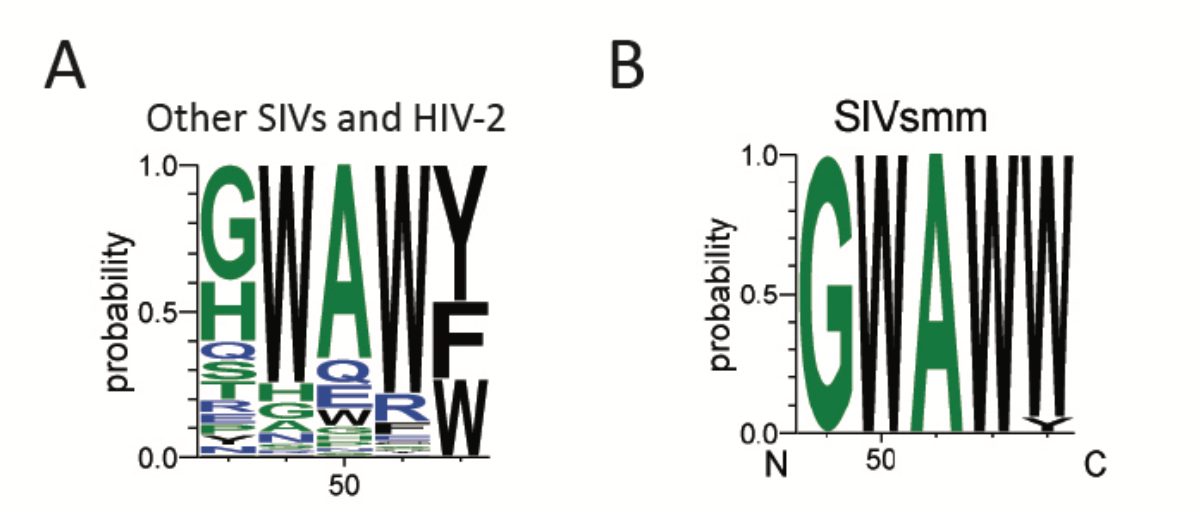
Sequence conservation of Vif loop 3. **A-B)** Logo plot of sequence variation of Vif loop 3 **A)** across HIV-2 and all SIVs, not including SIVcpz and SIVgor, (134 sequences) and **B)** of SIVsmm (19 sequences).

**Figure S11.**
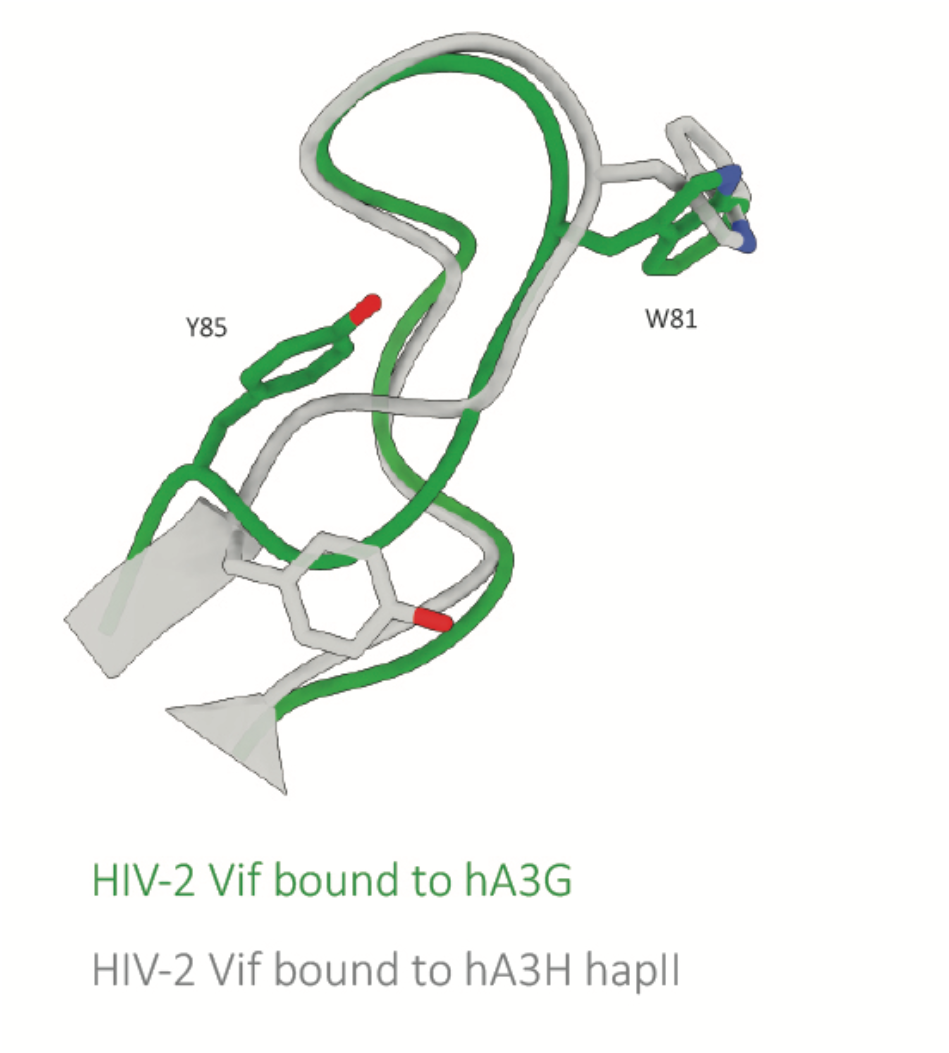
Structural comparison of HIV-2 Vif loop 5 when bound to hA3G or hA3H hap II. Alignment of HIV-2 Vif loop 5 when bound to hA3G (green) or hA3H hap II (grey; PDB: 9ZY0), with sidechains Y85 and W81 shown as sticks.

**Table S1.**
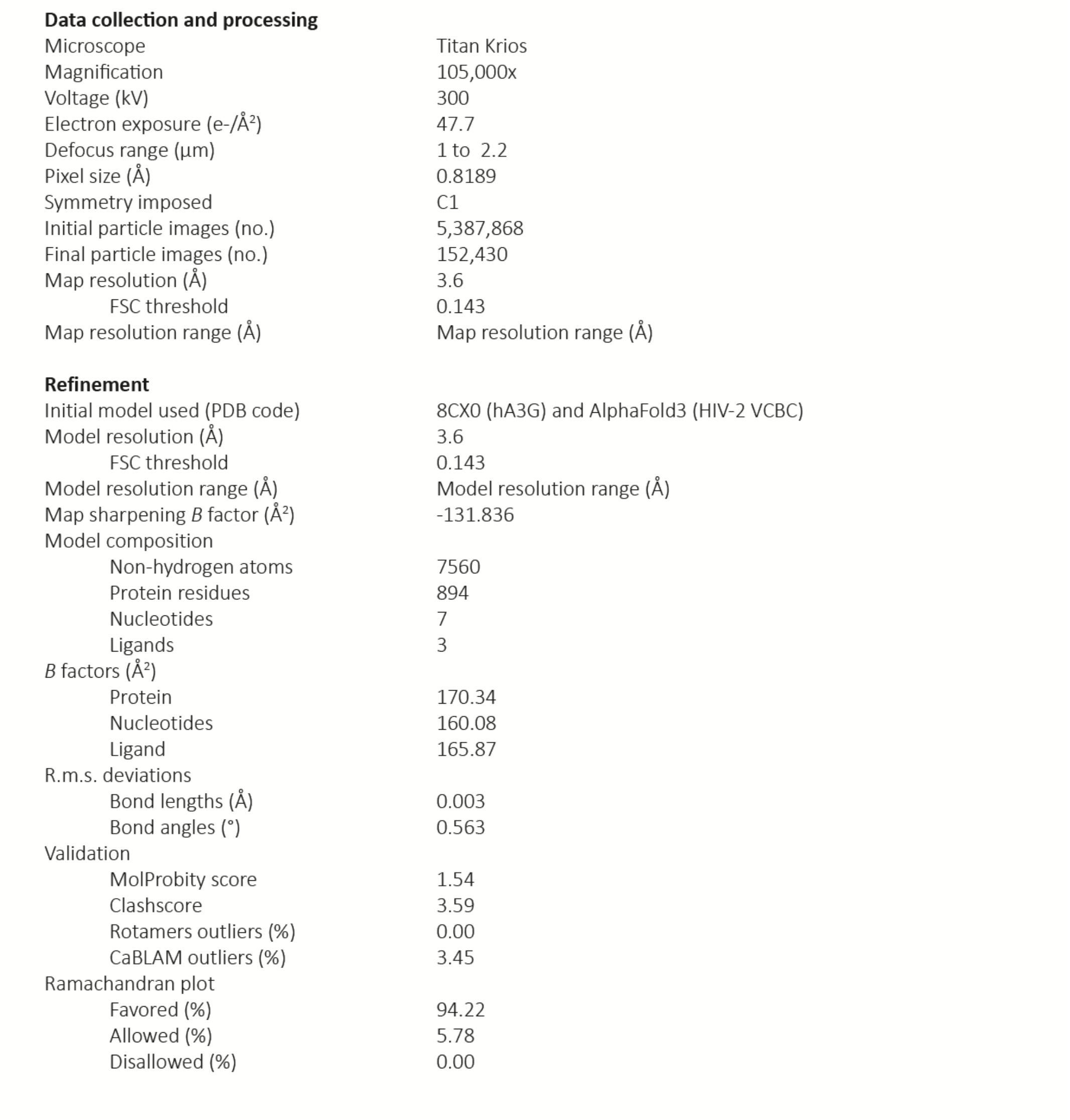
CryoEM data collection, refinement, and validation statistics.

**Data collection and processing**
|  |  |
| --- | --- |
| Microscope | Titan Krios |
| Magnification | 105,000x |
| Voltage (kV) | 300 |
| Electron exposure (e-/Å<sup>2</sup>) | 47.7 |
| Defocus range (μm) | 1 to 2.2 |
| Pixel size (Å) | 0.8189 |
| Symmetry imposed | C1 |
| Initial particle images (no.) | 5,387,868 |
| Final particle images (no.) | 152,430 |
| Map resolution (Å) | 3.6 |
| FSC threshold | 0.143 |
| Map resolution range (Å) | Map resolution range (Å) |
| **Refinement** |  |
| Initial model used (PDB code) | 8CX0 (hA3G) and AlphaFold3 (HIV-2 VCBC) |
| Model resolution (Å) | 3.6 |
| FSC threshold | 0.143 |
| Model resolution range (Å) | Model resolution range (Å) |
| Map sharpening *B* factor (Å<sup>2</sup>) | -131.836 |
| Model composition |  |
| Non-hydrogen atoms | 7560 |
| Protein residues | 894 |
| Nucleotides | 7 |
| Ligands | 3 |
| *B* factors (Å<sup>2</sup>) |  |
| Protein | 170.34 |
| Nucleotides | 160.08 |
| Ligand | 165.87 |
| R.m.s. deviations |  |
| Bond lengths (Å) | 0.003 |
| Bond angles (°) | 0.563 |
| Validation |  |
| MolProbity score | 1.54 |
| Clashscore | 3.59 |
| Rotamers outliers (%) | 0.00 |
| CaBLAM outliers (%) | 3.45 |
| Ramachandran plot |  |
| Favored (%) | 94.22 |
| Allowed (%) | 5.78 |
| Disallowed (%) | 0.00 |

**ST2.**
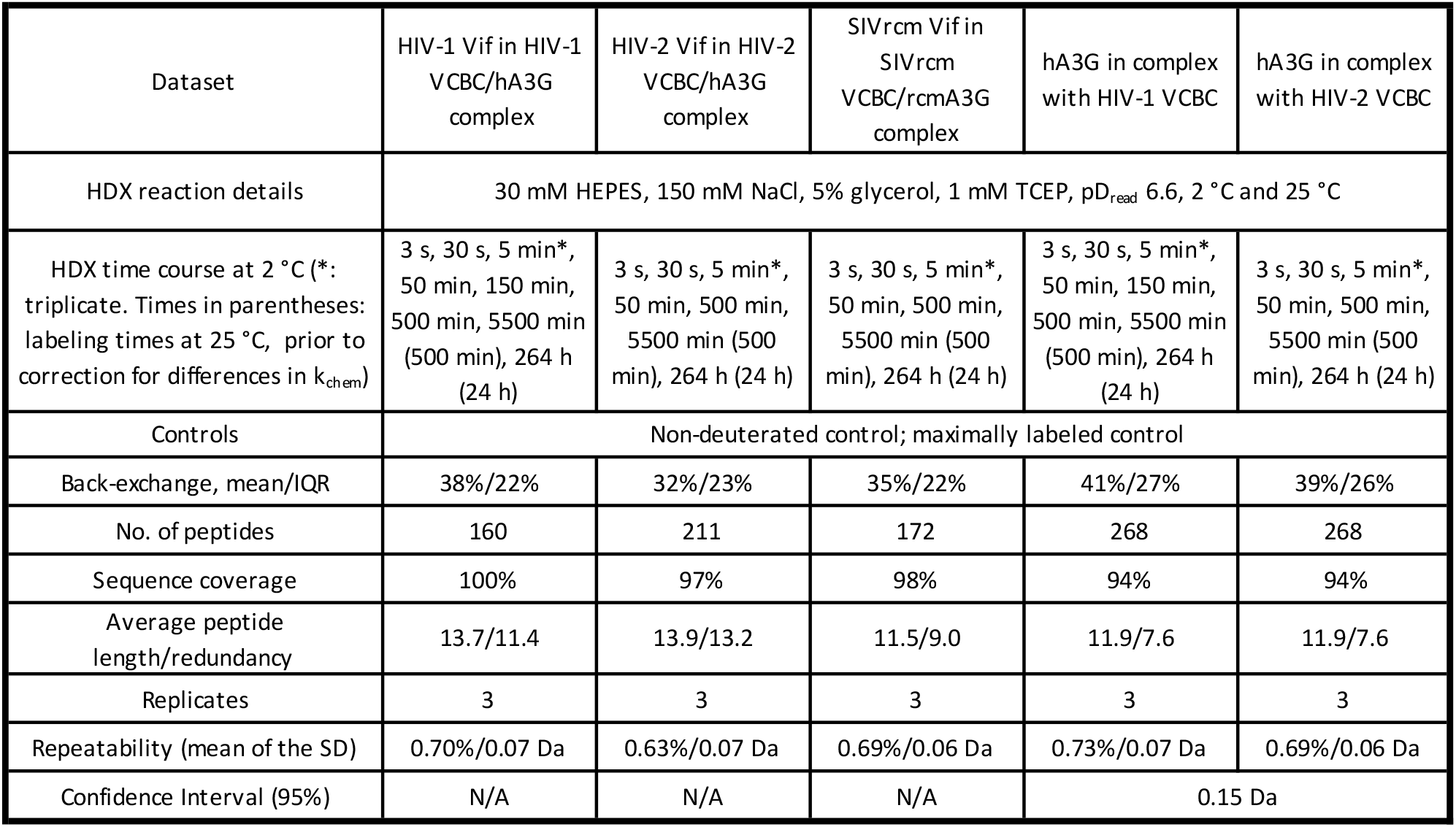
Biochemical and statistical details for HDX-MS.

ST2. Biochemical and statistical details for HDX-MS
| Dataset | HIV-1 Vif in HIV-1 VCBC/hA3G complex | HIV-2 Vif in HIV-2 VCBC/hA3G complex | SIVrcm Vif in SIVrcm VCBC/rcmA3G complex | hA3G in complex with HIV-1 VCBC | hA3G in complex with HIV-2 VCBC |
| --- | --- | --- | --- | --- | --- |
| HDX reaction details | 30 mM HEPES, 150 mM NaCl, 5% glycerol, 1 mM TCEP, $pD_{read}$ 6.6, 2 °C and 25 °C | | | | |
| HDX time course at 2 °C (*: triplicate. Times in parentheses: labeling times at 25 °C, prior to correction for differences in $k_{chem}$) | 3 s, 30 s, 5 min*, 50 min, 150 min, 500 min, 5500 min (500 min), 264 h (24 h) | 3 s, 30 s, 5 min*, 50 min, 500 min, 5500 min (500 min), 264 h (24 h) | 3 s, 30 s, 5 min*, 50 min, 500 min, 5500 min (500 min), 264 h (24 h) | 3 s, 30 s, 5 min*, 50 min, 150 min, 500 min, 5500 min (500 min), 264 h (24 h) | 3 s, 30 s, 5 min*, 50 min, 500 min, 5500 min (500 min), 264 h (24 h) |
| Controls | Non-deuterated control; maximally labeled control |  |  |  |  |
| Back-exchange, mean/IQR | 38%/22% | 32%/23% | 35%/22% | 41%/27% | 39%/26% |
| No. of peptides | 160 | 211 | 172 | 268 | 268 |
| Sequence coverage | 100% | 97% | 98% | 94% | 94% |
| Average peptide length/redundancy | 13.7/11.4 | 13.9/13.2 | 11.5/9.0 | 11.9/7.6 | 11.9/7.6 |
| Replicates | 3 | 3 | 3 | 3 | 3 |
| Repeatability (mean of the SD) | 0.70%/0.07 Da | 0.63%/0.07 Da | 0.69%/0.06 Da | 0.73%/0.07 Da | 0.69%/0.06 Da |
| Confidence Interval (95%) | N/A | N/A | N/A | 0.15 Da |  |

